# Immune cell type signature discovery and random forest classification for analysis of single cell gene expression datasets

**DOI:** 10.1101/2023.03.24.534078

**Authors:** Bogac Aybey, Sheng Zhao, Benedikt Brors, Eike Staub

**Affiliations:** Merck Healthcare KGaA, Clinical Measurement Sciences, Oncology Bioinformatics, Darmstadt, Germany; Faculty of Biosciences, Heidelberg University, Heidelberg, Germany; Division of Applied Bioinformatics, German Cancer Research Center (DKFZ), Heidelberg, Germany; German Cancer Consortium (DKTK), German Cancer Research Center (DKFZ) Heidelberg, Germany

**Keywords:** single-cell RNA sequencing, gene signature discovery, cell type classification, machine learning, clustering, immunology

## Abstract

**Background:** Robust immune cell gene expression signatures are central to the analysis of single cell studies. Nearly all known sets of immune cell signatures have been derived by making use of only single gene expression datasets. Utilizing the power of multiple integrated datasets could lead to high-quality immune cell signatures which could be used as superior inputs to machine learning-based cell type classification approaches.

**Results:** We established a novel gene expression similarity-based workflow for the discovery of immune cell type signatures that leverages multiple datasets, here four single cell expression datasets from three different cancer types. We used our immune cell signatures to train random forest classifiers for immune cell type assignment of single-cell RNA-seq datasets. We obtained similar or better prediction results compared to commonly used methods for cell type assignment in two independent benchmarking datasets. Our gene signature set yields higher prediction scores than other published immune cell type gene sets in our random forest approach.

**Discussion and conclusion:** We demonstrated the quality of our immune cell signatures and their strong performance in a random forest-based cell typing approach. We argue that classifying cells based on our comparably slim sets of genes accompanied by a random forest-based approach not only matches or outperforms widely used published approaches. It also facilitates unbiased downstream statistical analyses of differential gene expression between cell types for 90% of all genes whose expression profiles have not been used for cell type classification.

## Introduction

Recent improvements in single cell technologies led to a multitude of singe cell RNA-seq (scRNA-seq) studies to understand the complex interplay of immune cells in human tissues (Zhu et al., 2017). The procedures to assign cell types to single cells in such data are a critical factor for the success of such studies (Lahnemann et al., 2020). There are hardly any ‘golden rules’ or generally accepted computational workflows for assigning cell type labels to cells. Still, cell types are often annotated manually: after unsupervised clustering of all cells a manual assignment of cell types to clusters is performed by assessing expression patterns of author-selected marker genes (Kiselev et al., 2019; Zhao et al., 2020). Manual cell type assignment after assessment of small sets of markers is error-prone: it can depend on habits and non-explicit rules and opinions of researchers. Both steps are major sources of irreproducibility in the field (Gibson, 2022). Many researchers demonstrate that larger sets of genes, gene signatures with sometimes dozens of genes, can provide more reliable information for cell type classification since e.g., not all cells express even the most used literature marker genes of their corresponding cell types (Grabski and Irizarry, 2022). Therefore, different sets of immune cell type gene expression signatures have been proposed (Abbas et al., 2005; Angelova et al., 2015; Bindea et al., 2013; Charoentong et al., 2017; Lahnemann et al., 2020; Rooney et al., 2015). The majority of these have been derived by analyses of bulk tissue-based RNA-sequencing (RNA-seq) datasets, the older studies have used microarray-derived gene expression data. Most recent signature sets have been derived by scRNA-seq data (Magen et al., 2019; Zhang et al., 2019b; Zilionis et al., 2019). However, only single gene expression experiments have been analyzed to derive or apply these signature sets: immune cell type gene signatures based on the integration of evidence from multiple published scRNA-seq gene expression studies have not been described so far.

There is a specific group of workflows for cell type annotation that depends on clustering the cells in one of the initial steps of cell type labeling (Aran et al., 2019; Stuart et al., 2019). Such cell type annotation processes are often strongly affected by clustering results which heavily depend on the clustering parameters (Zhao et al., 2020). Clustering and cell type annotation heavily affect downstream analysis. When clustering and differential gene expression testing has used information from all genes for cell typing this leads to statistical biases (Lahnemann et al., 2020). It is obvious that pre-requisites for statistical analyses are violated when first information from all gene profiles is used for establishing groups of cells, and later these same genes and groups are used for statistical tests to determine differential gene expression between groups (Gibson, 2022). Similarly, a violation of statistical independence can be observed in most automatic cell type annotation approaches which make use of a complete reference dataset plus the data to be annotated. The problem arises when the focus is set on highly variable genes in both, the reference, and new dataset, instead of only the reference data as information about gene variance in the new data is already used for re-training of the classification model (Pasquini et al., 2021). Such gene selection bias should be avoided in all cell type annotation workflows. Automated deterministic approaches that lead to reproducible cell type annotations would not only be beneficial for the interpretation of single studies, but they would also facilitate cross-comparison between single cell gene expression studies. Models obtained from pre-trained machine learning models that make use of only expression information of a small fraction of the genes and leave the majority of gene expression information untouched for unbiased downstream statistical analyses could be a significant step forward.

In our study, we established a novel discovery workflow to identify immune cell-specific gene expression signatures by leveraging multiple scRNA-seq tumor microenvironment datasets. Furthermore, to eliminate the sources of analytical bias and increase reproducibility, we utilized a random forest-based cell type classification operating on small sets of genes of our immune cell type signatures. We extensively tested the performance of our gene sets and our classification approach against other widely used cell annotation approaches on two peripheral blood mononuclear benchmarking datasets.

## Data and Methods

### Single cell RNA-seq datasets and quality control for genes and cells

We list all datasets used in this study for expression signature discovery, classifier training and validation in Tab. 1, along with quality control and normalization information. For the cancer datasets, we used only the tumor samples for our analyses. For all datasets, we removed cells expressing less than 200 genes and genes expressed in less than three cells. We used the cell type annotations published by the original authors for the purpose of final annotation of our signatures, comparing method and gene set performance.

**Tab. 1:**
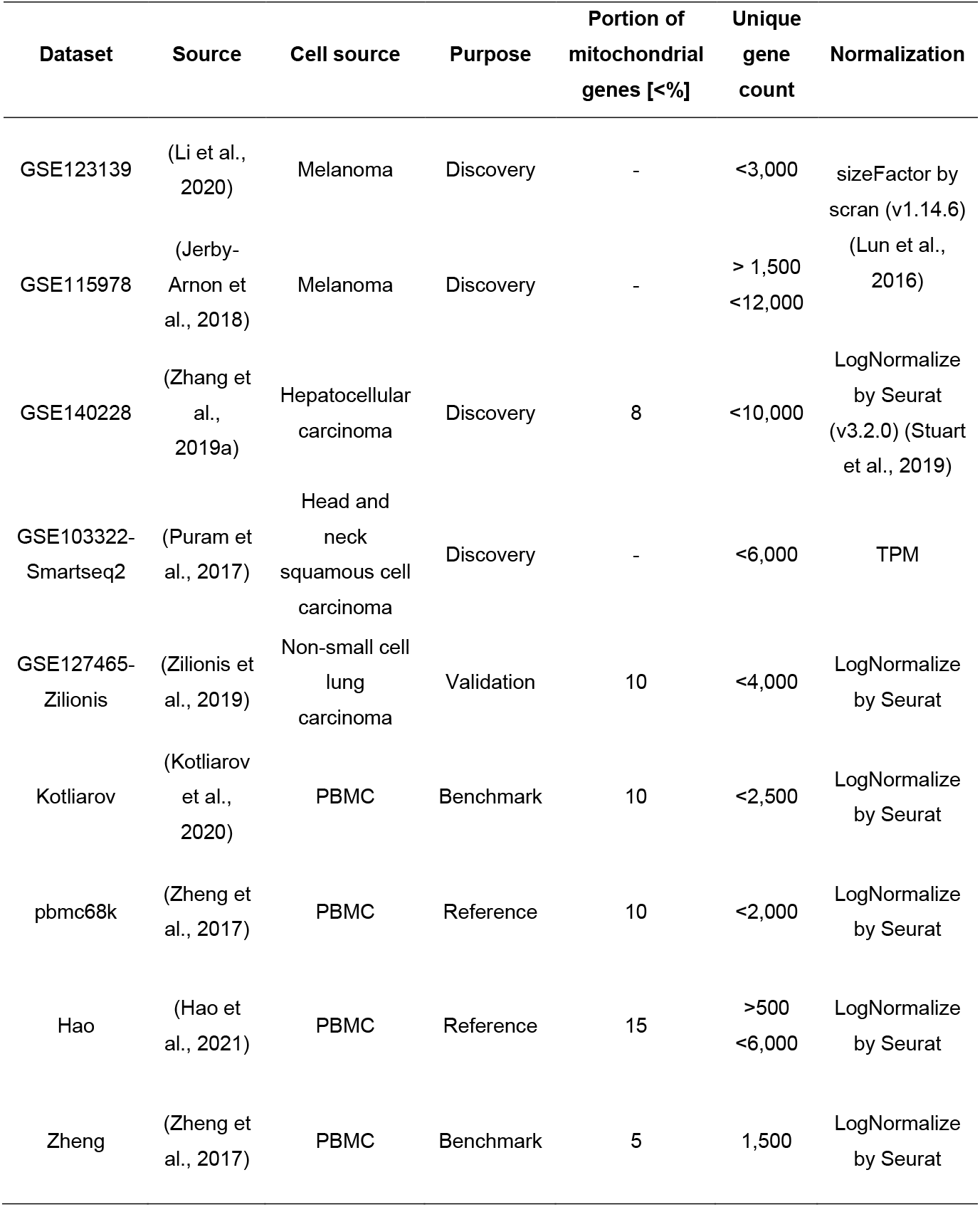
List of datasets used in this study along with their quality control measures.

### Gene sets and gene signatures for comparison to our approach

In addition to our own immune cell signatures, we investigated the following public immune cell signature repertoires: Abbas (Abbas et al., 2005), Charoentong (Charoentong et al., 2017), Angelova (Angelova et al., 2015), Becht (Becht et al., 2016), Bindea (Bindea et al., 2013), Newman (Newman et al., 2015), Nirmal (Nirmal et al., 2018) and Nieto (Nieto et al., 2021). For comparing our gene sets with others in a random forest approach, we used following cell types of the following published gene sets: a) Abbas: B cells, DCs, monocytes, NK, and T cells; b) Charoentong: B cells (general and memory), DCs (immature and pDC), monocytes, NK and CD4^+^ (regulatory, effector memory, central memory, and general) and CD8^+^ T cells (effector memory, central memory, and general); c) Angelova: B cells (memory and immature), DCs (pDC, immature, mDC and general), monocytes, NK, and CD4^+^ (regulatory, effector memory, and central memory) and CD8^+^ T cells (effector memory and central memory); d) Nieto: B and plasma cells, DCs (mDC, cDC and pDC), monocytes, NK cells, naïve T cells, CD4^+^ T (effector memory, transitional memory, memory/naïve and regulatory) and CD8^+^ T (effector memory and cytotoxic) cells.

### Exploratory analyses through dataset integration, dimension reduction, clustering, and feature selection

To distill robust immune cell type signatures and subsequently utilize those in cell type labeling tasks, we decided to leverage multiple scRNA-seq datasets from our discovery cohort. For the integration of multiple scRNA-seq datasets we used reciprocal principal component analysis (RPCA)-based integration from *Seurat* (v3.2.0) (Stuart et al., 2019). As suggested by the Seurat authors, for the integration of multiple large datasets we used the discovery dataset with the largest sample size, dataset GSE115978, as reference (option ‘reference-based RCPA’) which improves memory usage and run times. We selected 10k common highly variable genes (HVGs) as integration features. To find the anchors between dataset and the reference and to integrate the datasets, we used Seurat standard functions. As a final integrated dataset for our study, we obtained the corrected integrated expression matrix of 10k HVGs and 58,170 cells which we used as input to our immune cell signature discovery workflow.

To cluster genes with similar expression profiles in our integrated expression matrix, we used a density-based clustering approach. Prior to clustering we reduced the dimensionality of the integrated expression matrix using UMAP from *uwot* (v0.1.8) (Melville, 2019) on the gene dimension. Then, we performed density-based clustering using *dbscan* (v1.1.5) (Hahsler et al., 2019). We chose an optimal epsilon of 0.075 after plotting k-nearest neighbor distances in ascending order and analyzing the ‘knee’ point where maximum curvature was observed for a given minimum points (n=3) (minPts).

To refine the gene clusters, we calculated silhouette scores for individual genes and clusters using *cluster* (v2.1.0) (Maechler et al., 2021). As inputs we used the cluster labels from *dbscan* and the gene-by-gene distance matrix. We used as distance the so-called correlation distance for genes x and y: dx,y = 1 -*r*(x,y), where *r*(x,y) represents the Pearson correlation calculated from expression profiles of genes x and y.

To be able to evaluate the expression strength of each signature in each cell type, we calculated mean signature relative expression score for each cell and signature by averaging the Z-scaled (mean-centered and standardized across cells) expression values of all genes for a given signature (‘Average Z-Score method’).

For evaluation of the gene content of our signatures, we performed Gene Ontology (GO) analysis using *gprofiler2* (v0.1.9) (Kolberg and Raudvere, 2020) including correction for multiple testing. We measured the similarity of the gene sets (overlap of sets) by calculating the Jaccard index using *bayesbio* (v1.0.0) (McKenzie, 2016) and the Szymkiewicz–Simpson coefficient (Vijaymeena and Kavitha, 2016) between ours and all published signatures.

### Classification and performance benchmarking

For our immune cell type classification approach, we applied our immune cell type gene signatures as features in a random forest approach. For building random forest models utilizing *randomForest* (v4.6-14) (Liaw and Wiener, 2002) we used a ratio of 67:33 for random sampling of training and test data. We used only common genes between reference and query datasets as features. Prior to the training, we harmonized original cell type annotations from training datasets at medium-depth level shown in Supp. Tab. 1. For assessment of the performance of other published gene signatures, we applied an analogous procedure. Further, we used two different cell type annotation tools with default parameters: *Seurat* (v3.2.0) (Stuart et al., 2019) and *singleR* (v1.0.6) (Aran et al., 2019). Prior to cell type prediction using Seurat, we applied the standard pipeline to the query and reference dataset including log-normalization, finding and scaling HVGs. The anchors for cell type label transfer were determined between reference and query and cell type labels were then transferred. For singleR, predictions were obtained by providing normalized query and reference dataset along with cell type labels from a reference dataset.

Prior to the predictions, we harmonized original cell type annotations from benchmarking datasets at medium-depth level shown in Supp. Tab. 1. To evaluate the performance of the cell type prediction algorithms, we used six statistical metrics: accuracy, specificity, sensitivity, negative predictive value (NPV), positive predictive value (PPV) and F1-score. We reported the mean of each statistical metric for each algorithm.

## Results

Our study is structured into three parts: First, we comprehensively analyzed gene expression similarities across different single cell expression datasets. We discovered and refined gene modules to finally obtain robust gene signatures for immune cell types. Second, we tested the utility of our robust gene sets as classification features in a random forest-based (RF) immune cell type classification approach. We compared our approach with the two most widely used methods on two independent benchmarking datasets. Third, we examined how our gene sets are compared to other published gene signature sets or feature selection approaches, and how the way of choosing genes as features for classification influences the performance of a RF-based classification approach for cell type annotation.

### Discovery of robust immune cell type gene expression signatures by leveraging multiple scRNA-seq datasets

To discover robust immune cell signatures, we established an integrated density-based clustering workflow for scRNA-seq data (Fig. 1A). We integrated four scRNA-seq datasets of primary tumors from three cancer types (melanoma, hepatocellular carcinoma and head and neck squamous cell carcinoma). For the integration, we selected top 10k genes which were repeatedly variable across discovery datasets using *Seurat* (v3.2.0) where we applied reciprocal principal component-analysis (RPCA) based integration. We investigated the results of dataset harmonization by dimension reduction for the cell space of our integrated single cell gene expression matrix using the UMAP algorithm (Fig. 2A-B). We observed that cells were visually falling apart into separate clusters. The clustering was governed by cell type, not by study. Cells such as B cells, T cells or plasma cells clustered together despite coming from different studies. As expected, cell types uniquely measured in distinct studies, such as endothelial cells and fibroblasts, formed distinct clusters. The successfully integrated gene expression matrix comprised expression profiles of 10k genes for 58,170 cells from four datasets and subsequently served us to discover immune cell type-specific gene signatures.

**Fig. 1.**
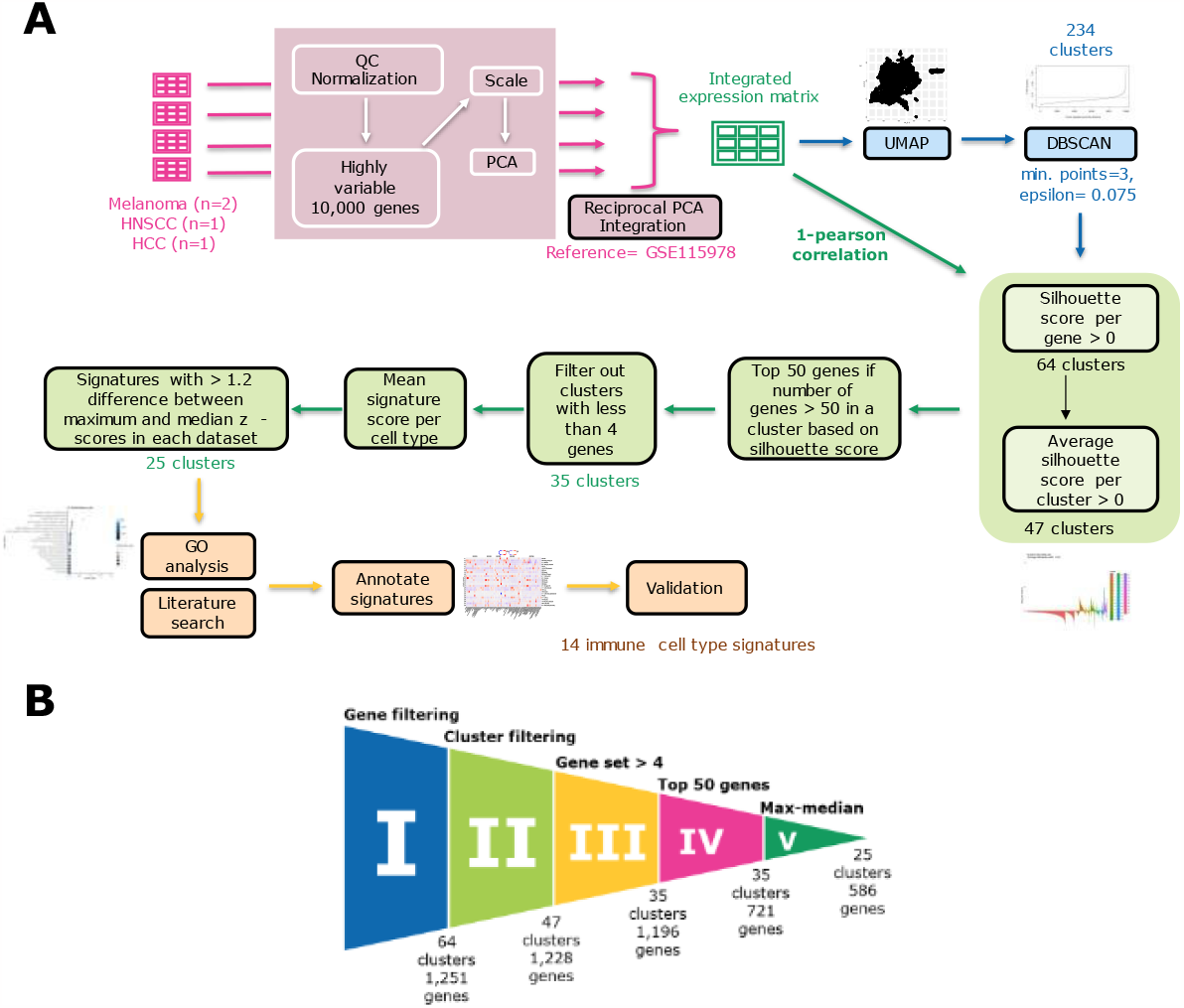
Immune cell type gene signature discovery workflow. **(A)** The workflow comprises the following steps: dataset integration, density-based clustering using DBSCAN, refinement of gene sets using filtering approaches based on silhouette scores and mean signature expression score, and annotating the signatures based on Gene Ontology analysis, literature search and mean signature expression score. **(B) Funnel plot showing the refinement process in each step**. The refinement process consists of five filtering steps: gene filtering, cluster filtering based on silhouette scores, selection of gene sets with minimum four genes, selection of top 50 genes and max-median filter based on mean signature expression scores. Each step is labeled from I to V. Final number of clusters and genes is shown after each filtering step.

**Fig. 2:**
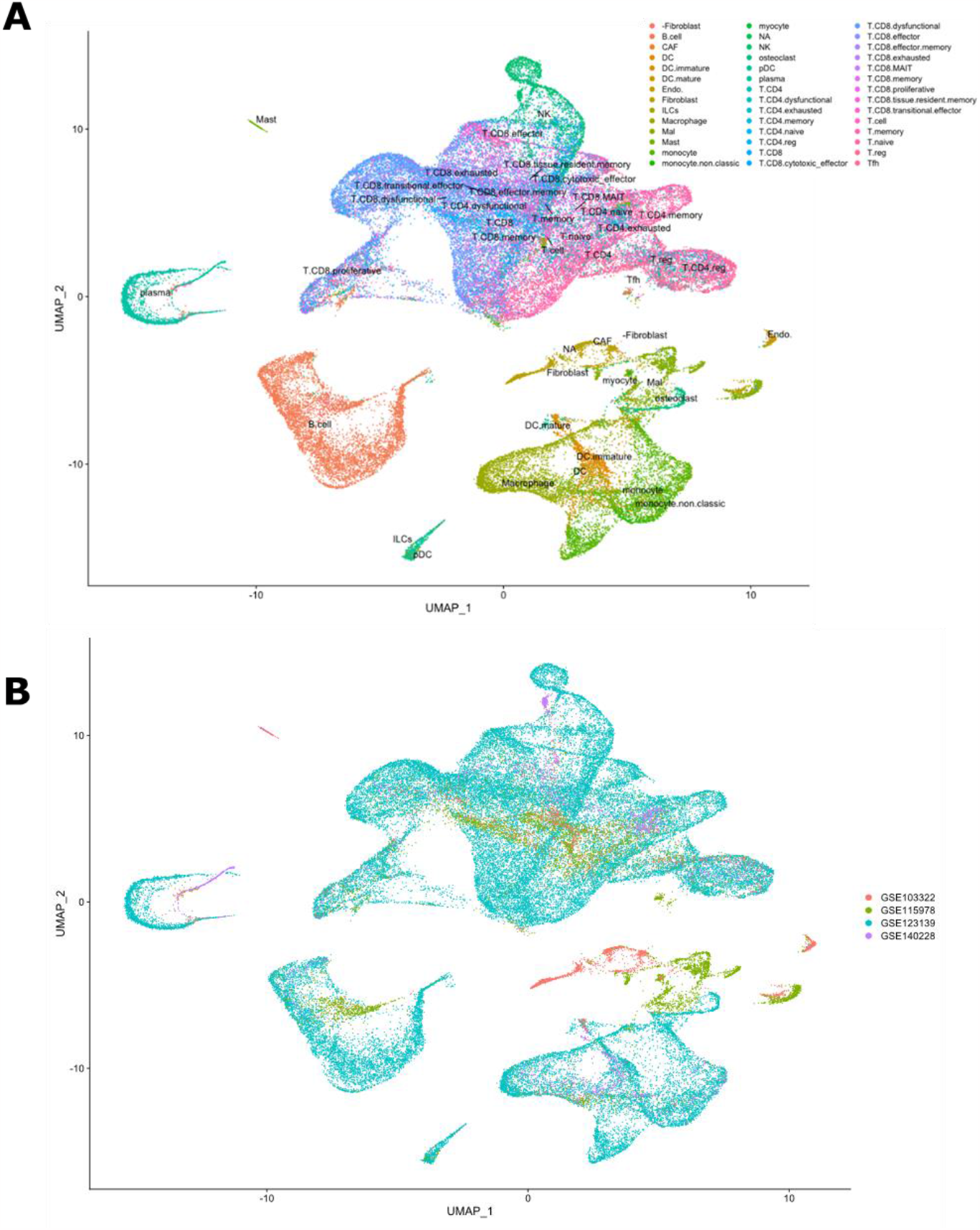
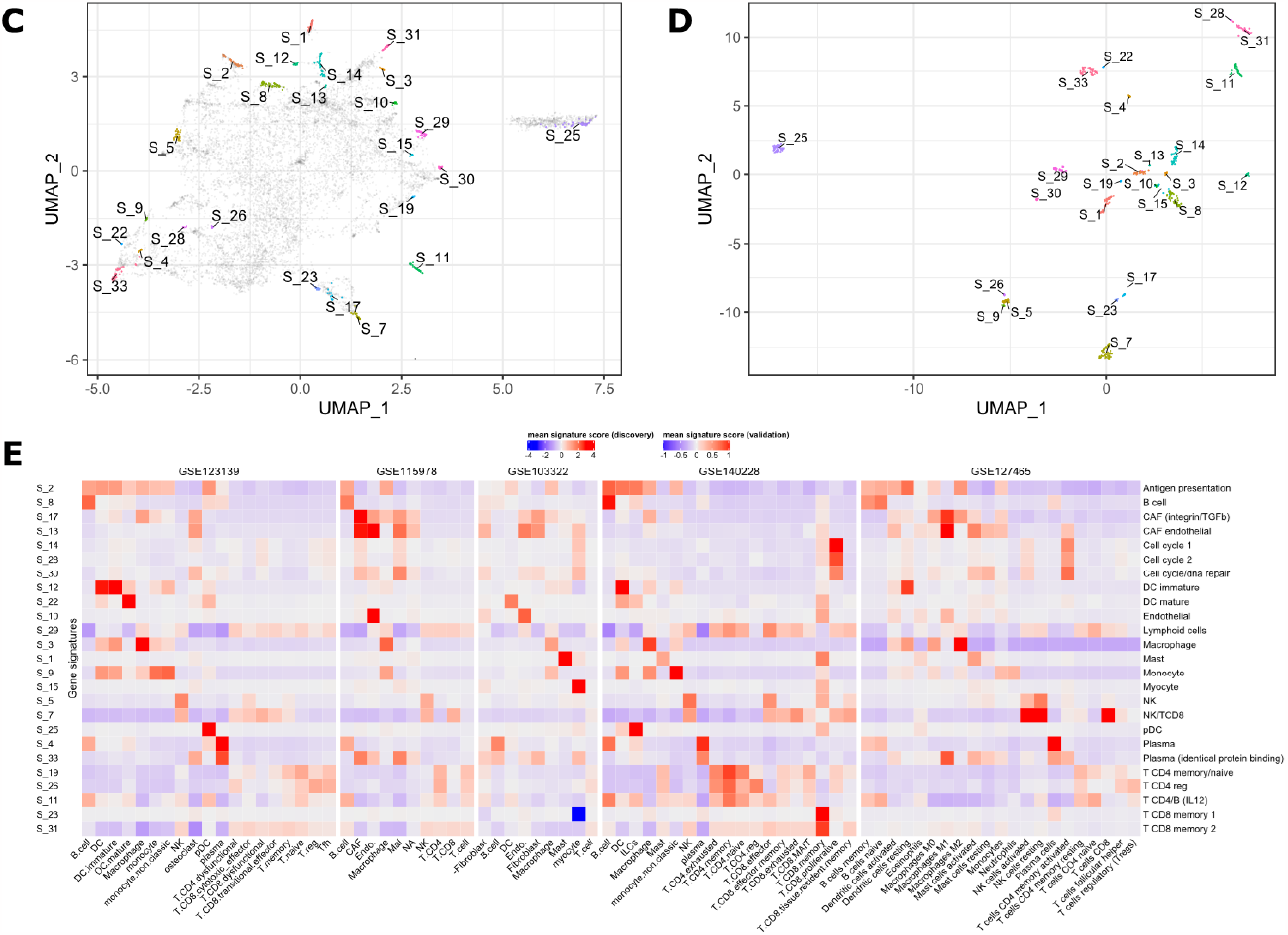
Integration, refinement process and results of our gene signatures. (A-B) UMAP plots of integrated expression matrix. Each point represents a single cell, and each cell is colored by cell type (A) or dataset (B). The cell type labels are taken from the original publications. Cell type labels are placed in the center of the cell type clusters. successful harmonization of the datasets can be seen. Abbreviations: CAF, cancer-associated fibroblast; DC, dendritic cell; ILCs, innate lymphoid cells, Mal, malignant cells; NK, natural killer; pDC, plasmacytoid dendritic cell; T MAIT, mucosal-associated invariant T cell; Tfh, T follicular helper, NA; unknown cell types. **(C-D) UMAP plots based on 10k HVGs or selected 586 genes from our gene signatures**. The dimension of the expression data of 10k common HVGs from integration step or our gene sets is reduced in the gene space. Each point represents a gene. UMAP1 and UMAP2 are plotted for each gene in x and y axis, respectively. In B and C genes in the final signatures are annotated in different colors whereas in B other genes are colored in gray. Cluster numbers are placed in the center of the clusters. Genes from each refined gene set collocate together with their corresponding gene set separated from other gene sets. **(E) Mean signature expression scores per cell type of refined gene signatures shown in the discovery and validation datasets**. Red and blue represent high and low mean signature expression scores, respectively. Rows represent the gene signature cluster numbers along with the manual annotations while columns represent the cell types defined by the original authors in the datasets. The signature annotation names contain cell type which the signature can detect and/or biological processes related to this signature and/or cell type.

To identify clusters or modules of similarly expressed genes, we developed a workflow that starts with dimensionality reduction for the gene space -not the cell space as used for the integration-followed by density-based clustering of 10k HVGs using the DBSCAN algorithm. We obtained 64 clusters of genes: initially, all genes of a cluster belonged to the cluster’s gene set. For the refinement of the gene content of each cluster, we utilized the silhouette scores (Fig 1B I-II). We calculated sequentially gene-and cluster-wise silhouette scores. First, we filtered out genes which did not fit to their clusters, i.e., those with negative silhouette score. Next, we filtered out clusters with negative average gene-wise silhouette scores since these were clusters with inherently incoherent expression profiles. Further, we filtered out clusters with less than four genes and only included the top 50 genes in each cluster based on the gene-wise silhouette scores (Fig. 1B III-IV). We assessed how strong the mean relative gene expression score (averaged Z scores of all genes in cluster) is in all cell types as they had been annotated in the original publications. We noticed that some gene clusters exhibit high expression signals across multiple cell types and decided to not further focus on such gene sets: we filtered out clusters when they exhibited only a small difference (<1.2 Z score units on natural log scale) between the cell type with maximum expression score and the cell type with median score in all discovery datasets (Fig. 1B V). We finally assessed the success of our gene selection strategy by performing a UMAP dimension reduction focusing on the gene space (Fig 2C-D). For each of the 25 gene sets we found that all genes within a set clustered to the exclusion of other genes not in a set, both for a UMAP plot of all 10k genes and for a UMAP re-analysis of all refined genes.

After gene set refinement, we finally obtained 25 gene sets that were subjected to a comprehensive annotation approach, i.e., a comparison with gene sets described in the original studies and gene sets from Gene Ontology (GO). For the former, we compared our gene sets with those used and/or discovered in the five original studies from which our discovery and validation data stem (Supp. Tab. 2). We assessed whether the genes in the clusters have been already mentioned as defining specific cell type(s) in the original studies. For endothelial cells, CD4 ^+^ T cells, B cells, CAF, CD4^+^ T regulatory cells, NK and CD8^+^ T cells along with cell cycle clusters most genes in those clusters have been already mentioned in at least one publication of at least one discovery dataset. This provided us with a preliminary basic annotation for our gene clusters. Next, we obtained the top 20 GO terms associated with each gene set to make use of the biological domain knowledge and to relate each signature to general or cell type-related biological processes (Supp. Fig. 1). The last type of information for our gene set annotations comes from an analysis of mean gene expression scores across all cell types (as identified by original authors) of our integrated dataset calculated using Average Z-Score method (Fig. 2E). After aligning these three types of information, we annotated the gene clusters in immune cell type annotations at a medium level of granularity (CD4^+^ T memory/naïve, CD4^+^ T regulatory, CD8^+^ T, CD8^+^ T memory, NK, pDC and DC, B cell, plasma cells, macrophages, monocytes and mast cells) such that a given cluster shows exclusive highest expression in a specific cell type in at least in one discovery dataset and/or matches with existing biological annotations. For 4 of 25 clusters, we observed high expression in multiple cell types: we annotated those gene sets with both cell type labels (NK/CD8^+^ T cell, B/CD4^+^ T cell, CAF/endothelial and lymphoid lineage). Conversely, if clusters cannot be assigned to a specific cell type but GO annotations clearly point to a biological process (antigen presentation, cell cycle and DNA damage) we annotated these 4 clusters with the biological process. We also obtained 4 non-immune cell type signatures (CAF, CAF/endothelial, endothelial and myocyte). Since we have aimed to discover immune cell type signatures, from now on we will only focus on 17 immune cell type signatures.

To validate our gene sets and the correctness of our immune cell type annotations, we used the scRNA-seq gene expression dataset GSE127465 as a validation dataset (Zilionis et al., 2019): it has not been used for discovery. We established mean signature expression scores for each cell using the Average Z-Score method and assessed these across cell types mentioned in the original study (Zilionis et al., 2019) (Fig. 2E). For two CD8^+^ T memory signatures (S_23 and S_31) and one plasma signature (S_33) we could not confirm their exclusive expression and we removed these three signatures from our signature repertoire. Of note, in our validation dataset since there were no pDC annotations, we could neither validate nor invalidate this signature but in this cluster, we had one of the most common pDC markers LILRA4 and thus keep this signature as valid. We could confirm that 14 out of 17 immune cell signatures showed exclusively high signature scores for their corresponding cell types and summarize in Tab. 2.

**Tab. 2:**
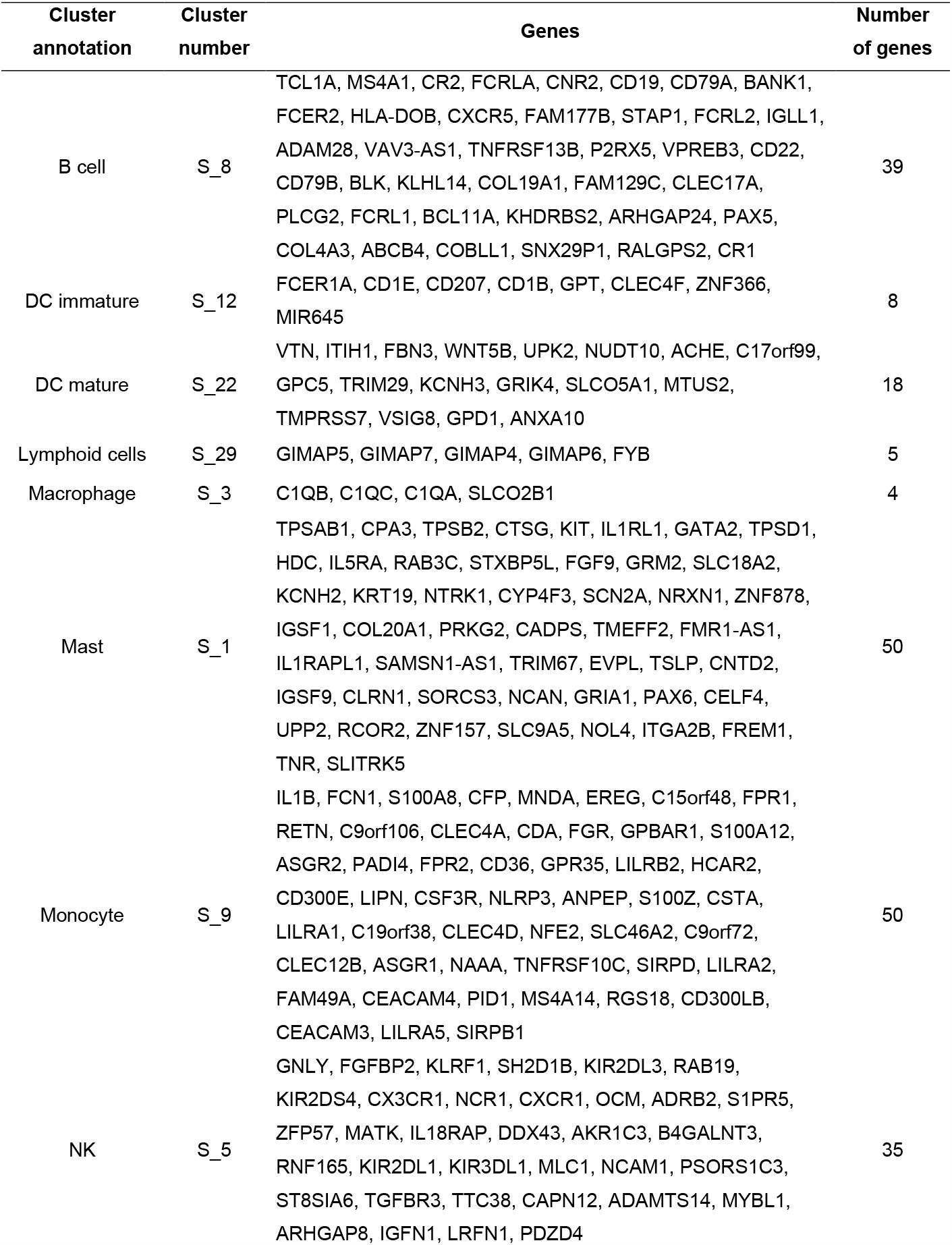
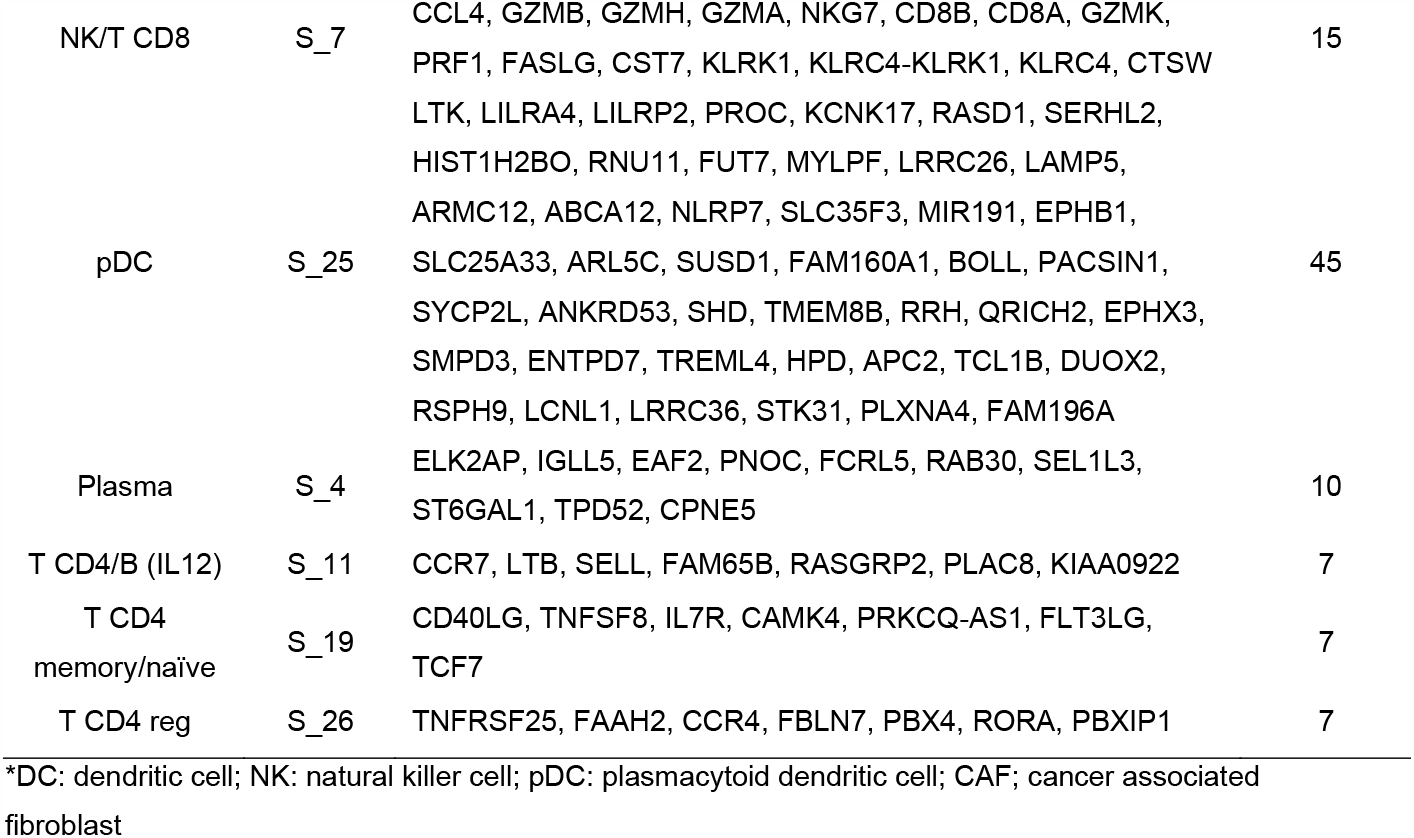
Summary of our refined immune cell type signatures.

It is possible that with our approach we just re-discovered gene sets that were already known. Thus, we compared our 14 immune cell type signatures with seven published immune cell signature repertoires using the Jaccard and Szymkiewicz-Simpson (S-S) indices. Our immune cell signatures had relatively low maximum Jaccard indices (0-0.31, median 0.09) (Supp. Fig. 2A). 50% of our immune cell signatures had maximum Jaccard indices below 0.1. The highest detected Jaccard indices were between our B cell signature and Nirmal, Newman and Becht B cell signatures (Jaccard idx= 0.31, 0.24 and 0.2, respectively), between our NK/CD8^+^ T and CD8 ^+^ T Newman signatures (Jaccard idx= 0.17) and between our monocyte and DC immature signatures and Newman monocyte and Becht DC/mDC signatures, respectively (Jaccard idx=0.17). Maximum S-S indices varied in a range from 0.06 to 0.89 (Supp. Fig. 2B). The maximum three S-S-indices were between our B cell and Becht B cell type signature (SSidx = 0.89), between our NK/CD8 ^+^ T and Newman CD8 ^+^ T signature (SSidx = 0.8), between our lymphoid signature and Nirmal T cell signature SSidx = 0.8) This shows that we have detected largely novel gene sets with significant differences in their composition compared to already published signatures, and that our signature gene sets were nearly always smaller than other gene sets.

In summary, we obtained 14 gene signatures for twelve distinct immune cell type populations, three distinct stromal cells and three distinct biological processes that exhibit high signature coherence in our four discovery datasets and one validation dataset.

### Using a limited number of cell type-specific genes combined with a random forest approach yields high prediction scores in comparison with commonly used algorithms

Most cell type annotation approaches use information from a very large number of HVGs (often 2k genes are used) for cell typing which might lead to problems with bias in down-stream statistical analyses where most important genes have been already used up for cell typing (Lahnemann et al., 2020). We hypothesized that using a small set of robust cell type signatures can predict as well as or better and can eliminate those biases. We applied our selected immune cell type genes (n= 209 genes) from our discovery workflow on a random forest cell type classification model: B cell (S_8), plasma cell (S_4), immature DC (S_12), pDC (S_25), monocyte (S_9), NK (S_5), NK/CD8^+^ T (S_7) and CD4^+^ T memory/naïve cell (S_19).

We compared our predictions with the most widely used cell typing tools, Seurat and singleR on the Kotliarov PBMC benchmarking dataset whose cell type annotation was based on cell surface marker protein expression which until now still the gold standard for assigning cell types (Kotliarov et al., 2020). First, we trained three reference datasets of different annotation quality or cellular environment: well-annotated PBMC-Hao (Hao et al., 2021), moderately-well annotated PBMC-pbmc68k (Zheng et al., 2017) and non-PBMC reference tumor-Zilionis (Zilionis et al., 2019). Hao reference yielded the best prediction rates among all methods on Kotliarov benchmarking dataset (Supp. Fig. 3A), and we proceeded with our comparison only based on Hao reference. Seurat used with 2k HVGs yielded the highest prediction scores followed by our approach which had high prediction scores (>0.9) in every metric (Fig. 3). We also executed Seurat using different number of HVGs. Using small number of HVGs (<500) did not perform well and had lower performance scores than our model in all six metrics (Supp. Fig. 3B). We note here that analyzing the test data for gene variability as in Seurat, followed by a new classifier training using the test data analysis results constitutes a violation of the machine learning principle to let no information from test data influence the classifier training and could lead to overoptimistic results. Our model showed better or similar results as two commonly used transfer methods and even better predictions than Seurat when using less HVGs.

**Fig. 3.**
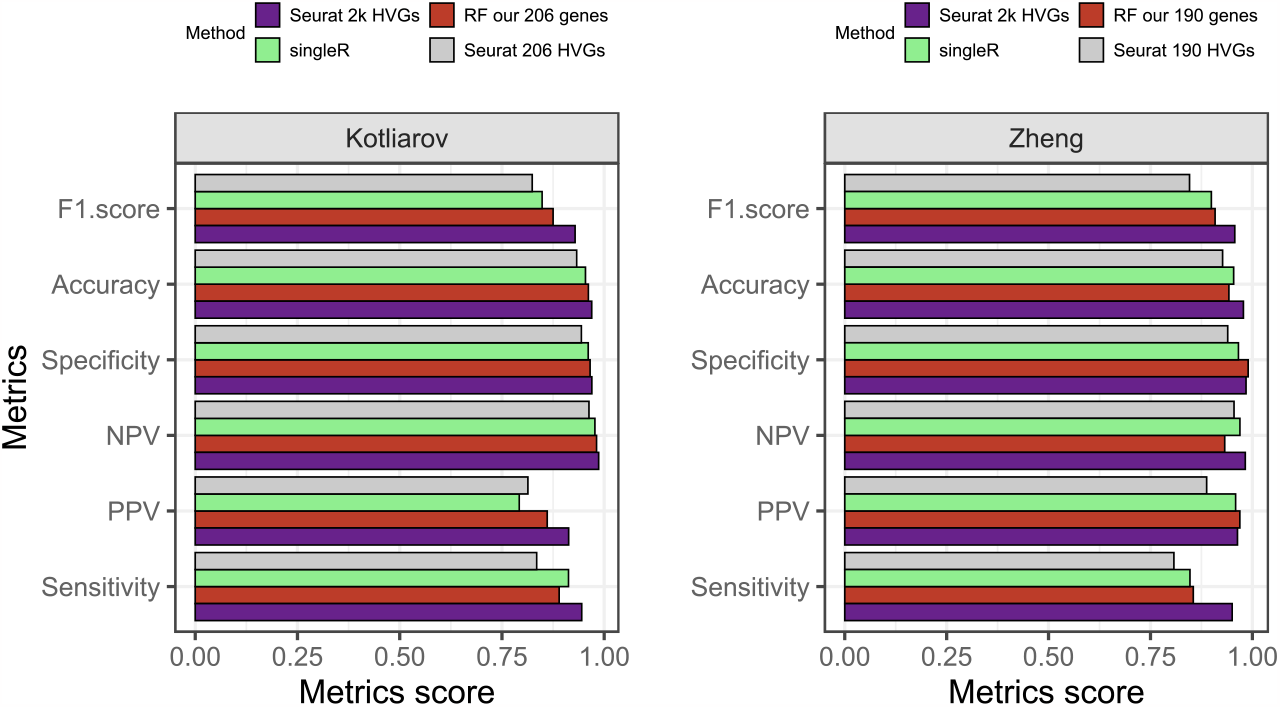
Our random forest model shows high prediction statistics compared to two commonly used tools in benchmarking datasets. Mean statistic metrics are displayed for each method. For reference dataset Hao dataset is used and as benchmarking dataset Kotliarov and Zheng datasets are used. Our model showed higher prediction scores compared to singleR and Seurat with similar number of genes as features.

We also evaluated all methods on a real ground truth dataset coming from FACS-sorted immune cells on Zheng dataset (Zheng et al., 2017). The dataset consists of nine immune cell populations: B cells, CD14 monocytes, naïve CD8^+^ T cells, cytotoxic CD8^+^ T cells, NK cells, memory CD4^+^ T cells, naïve CD4^+^ T cells, regulatory CD4^+^ T cells and helper CD4^+^ T cells. For the comparison, we downsampled 2k cells from each cell type and used the same set-up as previously described. Our model achieved high prediction scores (>0.85) in every metric except sensitivity (Fig. 3). Seurat using 2k HVGs had the highest prediction scores followed by our model while Seurat with same number of HVGs as the number of genes in our repertoire yielded the lowest prediction scores. Like for the Kotliarov benchmarking dataset, Seurat trained with small sets of genes (<500) performed worse than our model in all six metrics except NPV (Supp. Fig. 3C). So, our random forest approach based on a comparably small set of features obtained from our high-quality signature collection matched the performance or even outperformed the most widely used cell type annotation procedures in both, the Kotliarov dataset in which cell types had been annotated based on surface protein expression, and the Zheng dataset in which cells had been sorted by FACS.

### Using our immune cell type genes yields higher prediction than using other published immune cell type signature repertories

Finally, we assessed how our gene sets performed in a classification approach in comparison with four other published immune cell type signatures (hereafter referred to as the Angelova, Abbas, Charoentong and Nieto gene sets) (Abbas et al., 2005; Angelova et al., 2015; Charoentong et al., 2017; Nieto et al., 2021). We trained all random forest classifiers on subsets of genes (as defined by our or the four published signature sets) of the Hao dataset (Hao et al., 2021). We benchmarked the prediction results on scRNA-seq datasets of Kotliarov and Zheng (Kotliarov et al., 2020; Zheng et al., 2017). The comparator gene sets have been originally derived or previously utilized in different ways. The Nieto gene sets have been manually curated and have been applied on multiple scRNA-seq tumor microenvironment datasets; other three gene sets have been derived from bulk RNA-seq or microarray datasets. All four gene sets have been applied to study the tumor microenvironment. To highlight the lower bound of expected gene set performance we also assembled random sets of genes with the same number of genes which we had in our gene set repertoire (206 and 190 genes for Kotliarov and Zheng datasets, respectively).

The benchmarking results for all gene sets in our random forest classification approach with regard to overall prediction accuracy, F1 score, sensitivity, specificity, NPV, PPV are listed in Fig. 4. As expected, random genes showed the worst prediction scores. All gene sets clearly outperformed the randomly selected gene sets. Overall, the most favorable prediction results were obtained with our immune cell gene sets, followed by the Charoentong gene sets. Only for two benchmark parameters, sensitivity and NPV, on the Zheng dataset our gene sets were slightly outperformed or matching results from Charoentong, Nieto or Angelova gene sets. In summary, for both benchmarking datasets our random forest model trained on our comparably small sets of robust immune cell type marker genes overall showed superior performance when compared to the classifiers trained on four published gene repertoires.

**Fig. 4.**
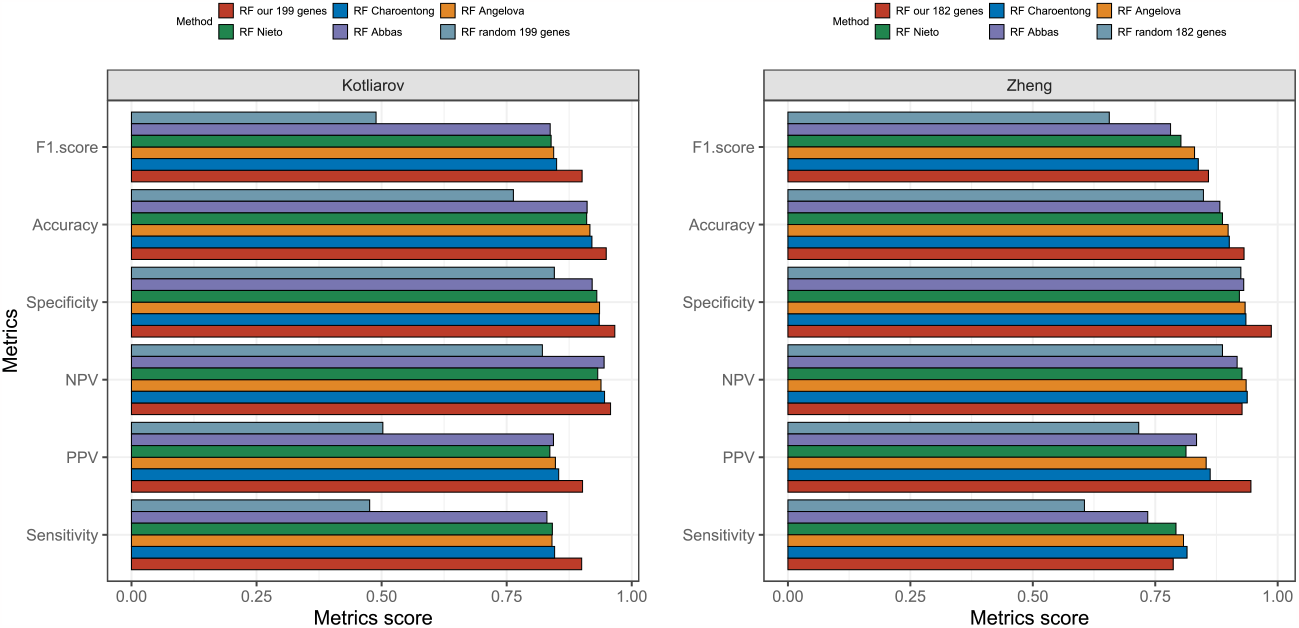
Comparison of our immune cell type signature repertoire with other published signatures on random forest approach in benchmarking datasets. Mean statistic metrics are shown for random forest models trained using our immune cell gene set and different published immune cell type repertoires (Abbas, Angelova, Charoentong and Nieto). We also include random forest model trained using same number of random genes as our immune cell type repertoire. Using our gene signatures shows higher prediction in benchmarking datasets, Kotliarov and Zheng datasets.

## Discussion and Conclusions

One main objective of our study was to discover and validate robust gene expression signatures for immune cell populations based on multiple single cell datasets. To do so, we established a novel workflow consisting of dataset integration, dimensionality reduction, density-based clustering, and cluster refinement methods. In our workflow we solely analyzed similarities between gene profiles across cells. In contrast to other approaches, we avoided selecting genes based on differential expression between cell type clusters, thus avoiding problems related to masking signals of small but important expression programs when focusing on the sample (here cell) level and not the gene level (Staub, 2012). Our benchmarking results show that our approach can yield small sets of marker genes with equal or better classification potential compared with published gene sets. Through application of our workflow on integrated single cell data from four studies we identified gene signatures for twelve distinct immune cell type populations. We detected a relatively low gene overlap (0-0.31 maximum Jaccard indices, 0.06-0.89 for maximum S-S indices) between our signatures and published signatures. This was expected since each published gene signature is derived using different analytical methods, different sequencing technologies, and different cell populations in the data that was used. Similarly, Nirmal et al. reported low concordance between their signatures and other published signatures (Nirmal et al., 2018). Interestingly, their B cell signature when compared to our B cell signature also yielded the highest Jaccard and SS indices. Low Jaccard scores to published signatures highlight the difficulty of the problem to identify robust signatures on the one hand, and the novelty of our approach to derive signatures on the other hand. Finally, they yield better cell type classification results compared to other published signatures. So, we argue that they are a reasonable choice as feature sets for cell type classification.

Even though our workflow robustly detected gene signatures for various cell types, we encountered the problem of separating NK and CD8^+^ T cells. We obtained one mixed NK/T CD8^+^ and one pure NK cell signature. The publications from which our data stem also reported the similarity between NK and CD8^+^ T cells on gene expression levels (Jerby-Arnon et al., 2018; Li et al., 2020; Zilionis et al., 2019). Similarly, for pbmc68k data Zheng et al. reported that cell assignment was not confident for NK and CD8^+^ T cells (Zheng et al., 2017). Considering the discussion on difficulty separating NK and CD8^+^ T cell populations in the literature, we argue that it is appropriate to use our two signatures in our random forest approach. The diagnostic separation of CD8 ^+^ T cells and NK cells by specific gene signatures should be subject of further studies.

Another aim of our study was to identify cell type classification approaches that are competitive when compared to existing methods for cell typing in complex scRNA-seq datasets. To this end, we developed a random forest classifier trained only on data from our gene sets. We compared our predictions with the two most used cell type annotation algorithms, Seurat and singleR. These were shown by others to perform well in different benchmarking setups, especially Seurat was gaining attention and currently is probably the most widely used cell type annotation tool (Huang et al., 2021). On our PBMC benchmarking datasets, Kotliarov and Zheng, we showed that our random forest model trained only on our gene signature collection achieved very high prediction scores that are close to the best performing Seurat approach. However, Seurat uses 10-fold more genes (2k HVGs instead of ∼200 distinct signature genes) and is not entirely unbiased since the classifier is trained on genes that were partially selected based on their high variance in the benchmark test set. The selection of HVGs might compromise those genes that are affected by important experimental variables such as treatment and time point. Especially for these genes it is important that results from downstream statistical analysis stay unbiased.

Some reports suggest that there are considerable differences in gene expression profiles of immune cells in different cellular environments, and that this leads to poor performance when an approach for cell typing is developed in one tissue context and then used in another tissue context (Nirmal et al., 2018; Pollara et al., 2017; Schelker et al., 2017). In our approach, we did not perceive tissue switching as a problem. We trained our classifiers on tumor microenvironment data and tested them on PBMC data with excellent performance. Furthermore, an important difference between other cell typing methods and our method is that we use only ∼200 genes (10% of 2k HVGs) for cell typing. For downstream analysis we therefore leave >90% of genes and their data untouched by classification. This means that >90% of genes can be subjected to downstream differential gene expression testing without bias. Such bias can arise when the same gene profiles are first used to establish cell type information (for example through clustering-based cell tagging) and later are tested for differential expression between groups that have already been defined using the same profiles. This is especially important for more complex experimental designs since most cell typing approaches face the problem of downstream analysis bias (Lahnemann et al., 2020). Currently, most single cell datasets do not have multiple experimental variables. In the future, single cell experimental set-ups could become more complex, especially when complex perturbations or time courses are to be analyzed. Then, providing cell types in a way that allows unbiased downstream analysis will become increasingly important.

In summary, our workflow is a valid approach to discover novel robust gene sets based on gene similarities from multiple scRNA-seq datasets. We demonstrated the superior performance of our immune cell type signatures compared to other gene sets in a random forest cell typing approach on two benchmarking datasets. In addition, we showed that random forest classifiers that use our genes sets as features match the performance of most widely used approaches for cell typing in scRNA-seq data, even though they do not make use of expression from 90% of genes. For these 90% of genes this generates opportunities for unbiased downstream statistical analyses making use of cell type information. Our results make us confident that our immune cell type signatures combined with a random forest approach can be used to analyze further complex single cell data. With even more data becoming available in a quickly growing universe of scRNA-seq gene expression data we are sure that our signature discovery approach can lead to many further cell type signature discoveries, also in other tissues than the tumor microenvironment or PBMCs.

## Declarations

### Authors’ contributions

**Eike Staub:** conceptualization, methodology, supervision, funding acquisition, reviewing, and editing. **Bogac Aybey:** conceptualization, methodology, software, visualization, data curation, formal analysis, investigation, writing, original draft preparation, reviewing, and editing. **Sheng Zhao:** data curation and supervision. **Benedikt Brors:** supervision and reviewing. All authors have read and agreed to the published version of the manuscript.

## Acknowledgments

We thank Dr. Felix Geist for valuable discussions and suggestions.

## Institutional Review Board Statement

We refer to the original authors for the institutional review board statement in publicly available datasets.

### Informed Consent Statement

We refer to the original authors for the informed consent in publicly available datasets.

### Data Availability Statement

All publicly available datasets used in this study have been listed in the methods section.

### Code availability

The scripts and outputs from this study will be deposited on https://github.com/bogacaybey/immune_signatures_discovery/.

