## Supplementary Figures and Tables for "Immune cell type signature discovery and random forest classification for analysis of single cell gene expression datasets": Supplementary Fig legends.docx

**Supp. Fig. 1: GO Analysis of final signatures.** Top 20 GO terms versus precision (intersection size / query size) are drawn for each signature. For multiple testing correction, false discovery rate is used with 0.05 as α threshold. Intersection size and p-values of each term are shown in the legends. The sizes of the intersection sizes are proportional to the number of intersection size while p-values are drawn in a scale from dark blue (high) to light blue (low).

**Supp. Fig. 2. Heatmap of Jaccard index scores (A) and Szymkiewicz–Simpson coefficients (B) between our immune cell type signatures and seven other published immune cell type signatures.** Jaccard index scores and Szymkiewicz–Simpson coefficients are calculated between our 14 refined gene signatures (rows) and seven published cell signatures (columns). The number of genes in each gene set has been indicated in brackets.

**Supp. Fig. 3. Prediction metrics plots. (A) Prediction metrics observed using different reference datasets.** Mean statistic metrics are displayed for a given reference dataset using different methods: our random forest model based on our immune cell type gene signatures, Seurat and singleR. For reference dataset Hao, pbmc68k and Zilionis datasets are tested and as benchmarking dataset Kotliarov dataset is used. **(B-C) Prediction metrics of Seurat with increasing HVGs.** In an interval of 100 HVGs, we executed Seurat using Hao reference dataset on Kotliarov (B) or Zheng (C) benchmarking dataset. We reported mean scores for six different statistic metrics.
