## Supplementary Figures and Tables for "Immune cell type signature discovery and random forest classification for analysis of single cell gene expression datasets": Supplementary Tables.docx

**Supp. Tab. 1. Medium-depth level cell type harmonization.** For benchmarking and reference datasets cell types are summarized into medium-level categories. Larger groups are in bold under benchmarking dataset and specific cell types belonging to those groups in every dataset are listed. Other cells category from the reference datasets are removed from the training.

| **Groups** | **Benchmarking datasets** | | **Reference datasets** | | |
| --- | --- | --- | --- | --- | --- |
|  | **Kotliarov** | **Zheng** | **Zillionis** | **Hao** | **pbmc68k** |
| **B** | B transitional  B switched  B unswitched | B | B memory  B naive  Plasma | B memory  B naive  B intermediate  Plasma | B |
| **Monocyte** | CD14 monocyte  CD16 monocyte | CD16 monocyte | CD14 monocyte  CD16 monocyte | CD14 monocyte  CD16 monocyte | Monocyte |
| **DC** | DC  pDC | - | DC activated  DC resting | DC  pDC | DC |
| **NK** | NK | NK | NK activated  NK resting | NK | NK |
| **T CD4** | T CD4 memory  T CD4 naïve | T CD4 naïve  T CD4 memory  T reg (T CD4 regulatory)  T CD4 helper | T CD4 memory activated  T CD4 memory resting  T CD4 naïve  Tfh (T CD4 follicular helper cells)  T reg | T CD4 naïve  T CD4 EM (effector memory)  T CD4 CM (central memory)  T CD4 activated  T reg  T CD4 CTL (cytotoxic activity)  T CD4 proliferating | T CD4 naïve  T CD4 memory  T reg  T CD4 Th2 |
| **T CD8** | T CD8 memory  T CD8 naïve | T CD8 cytotoxic  T CD8 naïve | T CD8 | T CD8 naïve  T CD8 EM  T CD8 CM  T CD8 activated | T CD8 naïve  T CD8 |
| **Other cells** | Double negative T cells (dn T), unconventional CD161^hi^/CD3^+^/CD8^+^ T cells, hematopoietic stem cells (HSC) | - | Eosinophils, macrophages, mast cells, neutrophils | Mucosal-associated invariant T cells, gamma delta T cells (gdT), dnT, HSC, innate lymphoid cells, erythrocytes, platelets | - |

**Supp. Tab. 2. Literature comparison of genes in our gene sets.** Genes in the final signatures are compared with the original publications. Each gene and its corresponding signature in each gene set from the original publication are shown. Genes are also shown along with their cluster numbers and our final annotations.

| **Genes** | **Cluster number** | **Jerby-Arnon_2018** | **Tirosh_2016** | **Li_2020** | **Zilionis_2019** | **Our annotation** |
| --- | --- | --- | --- | --- | --- | --- |
| TPSAB1 | S_1 | - | - | - | Mast | Mast |
| CPA3 | S_1 | - | - | - | Mast | Mast |
| TPSB2 | S_1 | - | - | - | Mast | Mast |
| CTSG | S_1 | - | - | - | Mast | Mast |
| KIT | S_1 | - | - | - | Mast | Mast |
| IL1RL1 | S_1 | - | - | - | Mast | Mast |
| GATA2 | S_1 | Endothelial | - | - | Mast | Mast |
| TPSD1 | S_1 | - | - | - | Mast | Mast |
| HDC | S_1 | - | - | - | Mast | Mast |
| IL5RA | S_1 | - | - | - | - | Mast |
| RAB3C | S_1 | - | - | - | Mast | Mast |
| STXBP5L | S_1 | - | - | - | - | Mast |
| FGF9 | S_1 | - | - | - | - | Mast |
| GRM2 | S_1 | - | - | - | - | Mast |
| SLC18A2 | S_1 | - | - | - | Mast | Mast |
| KCNH2 | S_1 | - | - | - | Mast | Mast |
| KRT19 | S_1 | - | - | - | Mast | Mast |
| NTRK1 | S_1 | - | - | - | Mast | Mast |
| CYP4F3 | S_1 | - | - | - | Neutrophil | Mast |
| SCN2A | S_1 | - | - | - | - | Mast |
| NRXN1 | S_1 | - | - | - | - | Mast |
| ZNF878 | S_1 | - | - | - | - | Mast |
| IGSF1 | S_1 | - | - | IG | - | Mast |
| COL20A1 | S_1 | - | - | - | - | Mast |
| PRKG2 | S_1 | - | - | - | Mast | Mast |
| CADPS | S_1 | - | - | - | Mast | Mast |
| TMEFF2 | S_1 | - | - | - | - | Mast |
| FMR1-AS1 | S_1 | - | - | - | - | Mast |
| IL1RAPL1 | S_1 | - | - | - | - | Mast |
| SAMSN1-AS1 | S_1 | - | - | - | - | Mast |
| TRIM67 | S_1 | - | - | - | - | Mast |
| EVPL | S_1 | - | - | - | Mast | Mast |
| TSLP | S_1 | - | - | - | - | Mast |
| CNTD2 | S_1 | - | - | - | - | Mast |
| IGSF9 | S_1 | - | - | IG | - | Mast |
| CLRN1 | S_1 | - | - | - | - | Mast |
| SORCS3 | S_1 | - | - | - | - | Mast |
| NCAN | S_1 | - | - | - | - | Mast |
| GRIA1 | S_1 | - | - | - | - | Mast |
| PAX6 | S_1 | - | - | - | - | Mast |
| CELF4 | S_1 | - | - | - | - | Mast |
| UPP2 | S_1 | - | - | - | - | Mast |
| RCOR2 | S_1 | - | - | - | - | Mast |
| ZNF157 | S_1 | - | - | - | - | Mast |
| SLC9A5 | S_1 | - | - | - | - | Mast |
| NOL4 | S_1 | - | - | - | - | Mast |
| ITGA2B | S_1 | - | - | - | Mast | Mast |
| FREM1 | S_1 | - | - | - | Mast | Mast |
| TNR | S_1 | - | - | - | B-cell | Mast |
| SLITRK5 | S_1 | - | - | - | - | Mast |
| CD74 | S_2 | - | - | - | - | Antigen presentation |
| HLA-DRA | S_2 | Macrophage | - | - | - | Antigen presentation |
| HLA-DMB | S_2 | Macrophage | - | - | - | Antigen presentation |
| NAPSB | S_2 | B-cell | - | - | - | Antigen presentation |
| CYSLTR1 | S_2 | - | - | - | - | Antigen presentation |
| CD72 | S_2 | - | - | - | - | Antigen presentation |
| PLD4 | S_2 | - | - | - | pDC | Antigen presentation |
| LAT2 | S_2 | - | - | - | Mast | Antigen presentation |
| BTK | S_2 | - | - | - | Mast | Antigen presentation |
| CD180 | S_2 | - | - | - | - | Antigen presentation |
| HLA-DMA | S_2 | Macrophage | - | - | - | Antigen presentation |
| HLA-DQA1 | S_2 | Macrophage | - | - | - | Antigen presentation |
| NCF1 | S_2 | - | - | - | Neutrophil | Antigen presentation |
| CD83 | S_2 | - | - | - | pDC | Antigen presentation |
| HLA-DRB1 | S_2 | Macrophage | - | - | - | Antigen presentation |
| HLA-DRB5 | S_2 | Macrophage | - | - | - | Antigen presentation |
| HLA-DPA1 | S_2 | Macrophage | - | - | - | Antigen presentation |
| CD40 | S_2 | - | - | - | B-cell | Antigen presentation |
| HLA-DQB1 | S_2 | Macrophage | - | - | - | Antigen presentation |
| HVCN1 | S_2 | B-cell | B-cell | - | pDC | Antigen presentation |
| FCGR2B | S_2 | - | - | - | - | Antigen presentation |
| IRF8 | S_2 | B-cell | B-cell | - | pDC | Antigen presentation |
| GAPT | S_2 | - | - | - | - | Antigen presentation |
| NCF1B | S_2 | - | - | - | - | Antigen presentation |
| HLA-DPB1 | S_2 | Macrophage | - | - | - | Antigen presentation |
| NCF1C | S_2 | B-cell | - | - | - | Antigen presentation |
| PIK3AP1 | S_2 | - | - | - | pDC | Antigen presentation |
| MARCH1 | S_2 | - | - | - | - | Antigen presentation |
| DAPP1 | S_2 | - | - | - | B-cell | Antigen presentation |
| NAPSA | S_2 | - | - | - | pDC | Antigen presentation |
| FGD2 | S_2 | - | - | - | - | Antigen presentation |
| HLA-DQB2 | S_2 | - | - | - | - | Antigen presentation |
| DENND3 | S_2 | - | - | - | Neutrophil | Antigen presentation |
| TBC1D9 | S_2 | - | - | - | - | Antigen presentation |
| HLA-DQA2 | S_2 | B-cell | - | - | - | Antigen presentation |
| KMO | S_2 | - | - | - | - | Antigen presentation |
| C1QB | S_3 | Macrophage | Macrophage | - | - | Macrophage |
| C1QC | S_3 | Macrophage | Macrophage | - | - | Macrophage |
| C1QA | S_3 | Macrophage | Macrophage | - | - | Macrophage |
| SLCO2B1 | S_3 | Macrophage | Macrophage | - | - | Macrophage |
| ELK2AP | S_4 | B-cell | - | - | - | Plasma |
| IGLL5 | S_4 | B-cell | - | IG | Plasma | Plasma |
| EAF2 | S_4 | - | - | - | Plasma | Plasma |
| PNOC | S_4 | B-cell | - | - | Plasma | Plasma |
| FCRL5 | S_4 | B-cell | - | - | Plasma | Plasma |
| RAB30 | S_4 | - | - | - | Plasma | Plasma |
| SEL1L3 | S_4 | B-cell | - | - | pDC | Plasma |
| ST6GAL1 | S_4 | B-cell | - | - | - | Plasma |
| TPD52 | S_4 | - | - | - | Plasma | Plasma |
| CPNE5 | S_4 | - | - | - | Plasma | Plasma |
| GNLY | S_5 | NK | - | - | NK | NK |
| FGFBP2 | S_5 | NK | - | - | NK | NK |
| KLRF1 | S_5 | NK | - | - | NK | NK |
| SH2D1B | S_5 | NK | - | - | NK | NK |
| KIR2DL3 | S_5 | NK | - | - | NK | NK |
| RAB19 | S_5 | - | - | - | - | NK |
| KIR2DS4 | S_5 | - | - | - | - | NK |
| CX3CR1 | S_5 | - | - | - | NK | NK |
| NCR1 | S_5 | NK | - | - | NK | NK |
| CXCR1 | S_5 | - | - | - | Neutrophil | NK |
| OCM | S_5 | - | - | - | - | NK |
| ADRB2 | S_5 | - | - | - | Mast | NK |
| S1PR5 | S_5 | - | - | - | NK | NK |
| ZFP57 | S_5 | - | - | - | - | NK |
| MATK | S_5 | NK | - | - | NK | NK |
| IL18RAP | S_5 | NK | - | - | Neutrophil | NK |
| DDX43 | S_5 | - | - | - | - | NK |
| AKR1C3 | S_5 | - | - | - | NK | NK |
| B4GALNT3 | S_5 | - | - | - | - | NK |
| RNF165 | S_5 | - | - | - | - | NK |
| KIR2DL1 | S_5 | - | - | - | NK | NK |
| KIR3DL1 | S_5 | - | - | - | NK | NK |
| MLC1 | S_5 | - | - | - | NK | NK |
| NCAM1 | S_5 | NK | - | - | NK | NK |
| PSORS1C3 | S_5 | - | - | - | - | NK |
| ST8SIA6 | S_5 | - | - | - | Mast | NK |
| TGFBR3 | S_5 | - | - | - | NK | NK |
| TTC38 | S_5 | - | - | - | NK | NK |
| CAPN12 | S_5 | - | - | - | NK | NK |
| ADAMTS14 | S_5 | - | - | - | - | NK |
| MYBL1 | S_5 | NK | - | - | NK | NK |
| ARHGAP8 | S_5 | - | - | - | - | NK |
| IGFN1 | S_5 | - | - | - | - | NK |
| LRFN1 | S_5 | - | - | - | Neutrophil | NK |
| PDZD4 | S_5 | - | - | - | NK | NK |
| CCL4 | S_7 | T CD8 cytotoxic | - | - | NK | NK/TCD8 |
| GZMB | S_7 | NK ,T CD8 cytotoxic | - | - | pDC | NK/TCD8 |
| GZMH | S_7 | T CD8 cytotoxic | - | - | NK | NK/TCD8 |
| GZMA | S_7 | T-cell | - | - | NK | NK/TCD8 |
| NKG7 | S_7 | NK | T- cell | - | NK | NK/TCD8 |
| CD8B | S_7 | T CD8 | - | - | T cell | NK/TCD8 |
| CD8A | S_7 | T CD8 ,T CD8 Exhausted | T- cell | - | T cell | NK/TCD8 |
| GZMK | S_7 | T CD8 | T- cell | - | T cell | NK/TCD8 |
| PRF1 | S_7 | NK ,T-cell ,T CD8 cytotoxic | T- cell | - | NK | NK/TCD8 |
| FASLG | S_7 | T CD8 cytotoxic | - | - | NK | NK/TCD8 |
| CST7 | S_7 | T-cell ,T CD8 cytotoxic | T- cell | - | NK | NK/TCD8 |
| KLRK1 | S_7 | - | T- cell | - | NK | NK/TCD8 |
| KLRC4-KLRK1 | S_7 | - | - | - | - | NK/TCD8 |
| KLRC4 | S_7 | - | - | - | NK | NK/TCD8 |
| CTSW | S_7 | NK | - | - | NK | NK/TCD8 |
| TCL1A | S_8 | B-cell | - | - | pDC | B cell |
| MS4A1 | S_8 | B-cell | B-cell | - | B-cell | B cell |
| CR2 | S_8 | B-cell | - | - | B-cell | B cell |
| FCRLA | S_8 | B-cell | B-cell | - | B-cell | B cell |
| CNR2 | S_8 | B-cell | - | - | B-cell | B cell |
| CD19 | S_8 | B-cell | B-cell | - | B-cell | B cell |
| CD79A | S_8 | B-cell | B-cell | - | - | B cell |
| BANK1 | S_8 | B-cell | B-cell | - | B-cell | B cell |
| FCER2 | S_8 | B-cell | - | - | B-cell | B cell |
| HLA-DOB | S_8 | B-cell | B-cell | - | B-cell | B cell |
| CXCR5 | S_8 | B-cell | B-cell | - | B-cell | B cell |
| FAM177B | S_8 | - | - | - | B-cell | B cell |
| STAP1 | S_8 | B-cell | B-cell | - | pDC | B cell |
| FCRL2 | S_8 | B-cell | - | - | B-cell | B cell |
| IGLL1 | S_8 | B-cell | - | IG | Plasma | B cell |
| ADAM28 | S_8 | B-cell | - | - | B-cell | B cell |
| VAV3-AS1 | S_8 | - | - | - | - | B cell |
| TNFRSF13B | S_8 | B-cell | - | - | B-cell | B cell |
| P2RX5 | S_8 | B-cell | B-cell | - | B-cell | B cell |
| VPREB3 | S_8 | B-cell | B-cell | - | B-cell | B cell |
| CD22 | S_8 | B-cell | B-cell | - | B-cell | B cell |
| CD79B | S_8 | B-cell | B-cell | - | B-cell | B cell |
| BLK | S_8 | B-cell | B-cell | - | B-cell | B cell |
| KLHL14 | S_8 | - | - | - | B-cell | B cell |
| COL19A1 | S_8 | B-cell | - | - | - | B cell |
| FAM129C | S_8 | B-cell | B-cell | - | pDC | B cell |
| CLEC17A | S_8 | B-cell | B-cell | - | - | B cell |
| PLCG2 | S_8 | - | B-cell | - | B-cell | B cell |
| FCRL1 | S_8 | B-cell | B-cell | - | B-cell | B cell |
| BCL11A | S_8 | B-cell | B-cell | - | pDC | B cell |
| KHDRBS2 | S_8 | - | - | - | - | B cell |
| ARHGAP24 | S_8 | - | - | - | B-cell | B cell |
| PAX5 | S_8 | B-cell | B-cell | - | B-cell | B cell |
| COL4A3 | S_8 | B-cell | - | - | B-cell | B cell |
| ABCB4 | S_8 | - | - | - | B-cell | B cell |
| COBLL1 | S_8 | - | - | - | - | B cell |
| SNX29P1 | S_8 | B-cell | - | - | - | B cell |
| RALGPS2 | S_8 | B-cell | B-cell | - | B-cell | B cell |
| CR1 | S_8 | - | - | - | Neutrophil | B cell |
| IL1B | S_9 | Macrophage | - | - | Neutrophil | Monocyte |
| FCN1 | S_9 | Macrophage | - | - | - | Monocyte |
| S100A8 | S_9 | Macrophage | - | - | Neutrophil | Monocyte |
| CFP | S_9 | Macrophage | - | - | Neutrophil | Monocyte |
| MNDA | S_9 | Macrophage | Macrophage | - | Neutrophil | Monocyte |
| EREG | S_9 | - | - | - | - | Monocyte |
| C15orf48 | S_9 | Macrophage | - | - | Neutrophil | Monocyte |
| FPR1 | S_9 | Macrophage | Macrophage | - | Neutrophil | Monocyte |
| RETN | S_9 | - | - | - | - | Monocyte |
| C9orf106 | S_9 | - | - | - | - | Monocyte |
| CLEC4A | S_9 | Macrophage | Macrophage | - | Neutrophil | Monocyte |
| CDA | S_9 | - | - | - | Neutrophil | Monocyte |
| FGR | S_9 | - | Macrophage | - | Neutrophil | Monocyte |
| GPBAR1 | S_9 | Macrophage | - | - | - | Monocyte |
| S100A12 | S_9 | - | - | - | Neutrophil | Monocyte |
| ASGR2 | S_9 | - | - | - | - | Monocyte |
| PADI4 | S_9 | - | - | - | Neutrophil | Monocyte |
| FPR2 | S_9 | Macrophage | - | - | Neutrophil | Monocyte |
| CD36 | S_9 | - | - | - | pDC | Monocyte |
| GPR35 | S_9 | - | - | - | - | Monocyte |
| LILRB2 | S_9 | Macrophage | Macrophage | - | - | Monocyte |
| HCAR2 | S_9 | Macrophage | - | - | Neutrophil | Monocyte |
| CD300E | S_9 | Macrophage | Macrophage | - | - | Monocyte |
| LIPN | S_9 | - | - | - | - | Monocyte |
| CSF3R | S_9 | Macrophage | Macrophage | - | Neutrophil | Monocyte |
| NLRP3 | S_9 | Macrophage | Macrophage | - | - | Monocyte |
| ANPEP | S_9 | - | - | - | - | Monocyte |
| S100Z | S_9 | - | - | - | - | Monocyte |
| CSTA | S_9 | Macrophage | Macrophage | - | - | Monocyte |
| LILRA1 | S_9 | Macrophage | Macrophage | - | - | Monocyte |
| C19orf38 | S_9 | Macrophage | - | - | Neutrophil | Monocyte |
| CLEC4D | S_9 | - | - | - | Neutrophil | Monocyte |
| NFE2 | S_9 | - | - | - | - | Monocyte |
| SLC46A2 | S_9 | - | - | - | - | Monocyte |
| C9orf72 | S_9 | Macrophage | - | - | Neutrophil | Monocyte |
| CLEC12B | S_9 | - | - | - | - | Monocyte |
| ASGR1 | S_9 | - | - | - | - | Monocyte |
| NAAA | S_9 | Macrophage | - | - | - | Monocyte |
| TNFRSF10C | S_9 | - | - | - | Neutrophil | Monocyte |
| SIRPD | S_9 | - | - | - | - | Monocyte |
| LILRA2 | S_9 | Macrophage | - | - | Neutrophil | Monocyte |
| FAM49A | S_9 | Macrophage | - | - | - | Monocyte |
| CEACAM4 | S_9 | - | - | - | - | Monocyte |
| PID1 | S_9 | - | - | - | - | Monocyte |
| MS4A14 | S_9 | - | - | - | - | Monocyte |
| RGS18 | S_9 | Macrophage | - | - | Neutrophil | Monocyte |
| CD300LB | S_9 | Macrophage | - | - | - | Monocyte |
| CEACAM3 | S_9 | - | - | - | Neutrophil | Monocyte |
| LILRA5 | S_9 | Macrophage | Macrophage | - | Neutrophil | Monocyte |
| SIRPB1 | S_9 | Macrophage | Macrophage | - | Neutrophil | Monocyte |
| PLVAP | S_10 | Endothelial | Endothelial | - | pDC | Endothelial |
| SELP | S_10 | Endothelial | - | - | - | Endothelial |
| MEOX1 | S_10 | Endothelial | - | - | - | Endothelial |
| VWF | S_10 | Endothelial | Endothelial | - | - | Endothelial |
| FCN3 | S_10 | - | - | - | - | Endothelial |
| EDN1 | S_10 | - | - | - | Neutrophil | Endothelial |
| CYYR1 | S_10 | Endothelial | - | - | pDC | Endothelial |
| LTF | S_10 | - | - | - | - | Endothelial |
| TM4SF18 | S_10 | Endothelial | Endothelial | - | - | Endothelial |
| PLEKHG5 | S_10 | - | - | - | - | Endothelial |
| STPG2 | S_10 | - | - | - | - | Endothelial |
| NPR1 | S_10 | Endothelial | - | - | Plasma | Endothelial |
| JAG2 | S_10 | Endothelial | - | - | NK | Endothelial |
| FAM107A | S_10 | Endothelial | - | - | - | Endothelial |
| TAL1 | S_10 | - | - | - | Mast | Endothelial |
| LRRC70 | S_10 | Endothelial | - | - | - | Endothelial |
| EMCN | S_10 | Endothelial | Endothelial | - | - | Endothelial |
| RSPO3 | S_10 | - | - | - | - | Endothelial |
| ROBO4 | S_10 | Endothelial | Endothelial | - | - | Endothelial |
| CCDC68 | S_10 | - | - | - | - | Endothelial |
| ARHGEF15 | S_10 | Endothelial | - | - | - | Endothelial |
| ERG | S_10 | Endothelial | Endothelial | - | Mast | Endothelial |
| GIPC2 | S_10 | Endothelial | - | - | - | Endothelial |
| CLDN15 | S_10 | Endothelial | - | - | - | Endothelial |
| RALGAPA2 | S_10 | Endothelial | - | - | - | Endothelial |
| INHBB | S_10 | - | - | - | - | Endothelial |
| FAM155A | S_10 | - | - | - | - | Endothelial |
| GIPC3 | S_10 | - | - | - | - | Endothelial |
| CCR7 | S_11 | T CD4 naive ,T CD8 naive | - | - | T cell | T CD4/B (IL12) |
| LTB | S_11 | T CD4 Treg | - | - | pDC | T CD4/B (IL12) |
| SELL | S_11 | B-cell ,T CD4 naive ,T CD8 naive | - | - | Neutrophil | T CD4/B (IL12) |
| FAM65B | S_11 | B-cell ,T CD4 naive | - | - | - | T CD4/B (IL12) |
| RASGRP2 | S_11 | - | - | - | B-cell | T CD4/B (IL12) |
| PLAC8 | S_11 | - | - | - | pDC | T CD4/B (IL12) |
| KIAA0922 | S_11 | - | - | - | - | T CD4/B (IL12) |
| LTK | S_25 | - | - | - | pDC | pDC |
| LILRA4 | S_25 | Macrophage | - | - | pDC | pDC |
| LILRP2 | S_25 | - | - | - | - | pDC |
| PROC | S_25 | - | - | - | - | pDC |
| KCNK17 | S_25 | - | - | - | - | pDC |
| RASD1 | S_25 | - | - | - | pDC | pDC |
| SERHL2 | S_25 | - | - | - | pDC | pDC |
| HIST1H2BO | S_25 | - | - | - | - | pDC |
| RNU11 | S_25 | - | - | - | - | pDC |
| FUT7 | S_25 | - | - | - | pDC | pDC |
| MYLPF | S_25 | - | - | - | - | pDC |
| LRRC26 | S_25 | - | - | - | pDC | pDC |
| LAMP5 | S_25 | - | - | - | pDC | pDC |
| ARMC12 | S_25 | - | - | - | - | pDC |
| ABCA12 | S_25 | - | - | - | - | pDC |
| NLRP7 | S_25 | - | - | - | - | pDC |
| SLC35F3 | S_25 | - | - | - | pDC | pDC |
| MIR191 | S_25 | - | - | - | - | pDC |
| EPHB1 | S_25 | - | - | - | pDC | pDC |
| SLC25A33 | S_25 | - | - | - | - | pDC |
| ARL5C | S_25 | - | - | - | - | pDC |
| SUSD1 | S_25 | - | - | - | pDC | pDC |
| FAM160A1 | S_25 | - | - | - | pDC | pDC |
| BOLL | S_25 | - | - | - | - | pDC |
| PACSIN1 | S_25 | - | - | - | pDC | pDC |
| SYCP2L | S_25 | - | - | - | - | pDC |
| ANKRD53 | S_25 | - | - | - | - | pDC |
| SHD | S_25 | - | - | - | pDC | pDC |
| TMEM8B | S_25 | - | - | - | pDC | pDC |
| RRH | S_25 | - | - | - | - | pDC |
| QRICH2 | S_25 | - | - | - | - | pDC |
| EPHX3 | S_25 | - | - | - | - | pDC |
| SMPD3 | S_25 | - | - | - | pDC | pDC |
| ENTPD7 | S_25 | - | - | - | pDC | pDC |
| TREML4 | S_25 | - | - | - | Neutrophil | pDC |
| HPD | S_25 | - | - | - | - | pDC |
| APC2 | S_25 | - | - | - | - | pDC |
| TCL1B | S_25 | - | - | - | - | pDC |
| DUOX2 | S_25 | - | - | - | - | pDC |
| RSPH9 | S_25 | - | - | - | - | pDC |
| LCNL1 | S_25 | - | - | - | - | pDC |
| LRRC36 | S_25 | - | - | - | - | pDC |
| STK31 | S_25 | - | - | - | - | pDC |
| PLXNA4 | S_25 | - | - | - | pDC | pDC |
| FAM196A | S_25 | - | - | - | - | pDC |
| FCER1A | S_12 | - | - | - | Mast | DC immature |
| CD1E | S_12 | - | - | - | - | DC immature |
| CD207 | S_12 | - | - | - | - | DC immature |
| CD1B | S_12 | - | - | - | - | DC immature |
| GPT | S_12 | - | - | - | - | DC immature |
| CLEC4F | S_12 | - | - | - | - | DC immature |
| ZNF366 | S_12 | Endothelial | - | - | - | DC immature |
| MIR645 | S_12 | - | - | - | - | DC immature |
| ZSCAN22 | S_31 | - | - | - | - | T CD8 memory 2 |
| SGSM1 | S_31 | - | - | - | - | T CD8 memory 2 |
| SPIRE2 | S_31 | - | - | - | - | T CD8 memory 2 |
| LINC00574 | S_31 | - | - | - | - | T CD8 memory 2 |
| MS4A10 | S_31 | - | - | - | - | T CD8 memory 2 |
| DLX6-AS1 | S_31 | - | - | - | - | T CD8 memory 2 |
| ZNF805 | S_31 | - | - | - | - | T CD8 memory 2 |
| LINC00672 | S_31 | - | - | - | - | T CD8 memory 2 |
| C1QTNF6 | S_31 | - | - | - | - | T CD8 memory 2 |
| VSTM4 | S_31 | T CD8 Exhausted | - | - | - | T CD8 memory 2 |
| TUBA3FP | S_31 | - | - | - | - | T CD8 memory 2 |
| CEP19 | S_31 | - | - | - | - | T CD8 memory 2 |
| LMOD3 | S_31 | - | - | - | - | T CD8 memory 2 |
| SMYD4 | S_31 | - | - | - | - | T CD8 memory 2 |
| MOG | S_31 | - | - | - | Mast | T CD8 memory 2 |
| NXN | S_31 | - | - | - | - | T CD8 memory 2 |
| SCD5 | S_31 | T CD8 Exhausted | - | - | - | T CD8 memory 2 |
| PHYHD1 | S_31 | - | - | - | - | T CD8 memory 2 |
| GP6 | S_31 | - | - | - | - | T CD8 memory 2 |
| C14orf105 | S_31 | - | - | - | - | T CD8 memory 2 |
| RAB36 | S_31 | - | - | - | - | T CD8 memory 2 |
| ZNF785 | S_31 | - | - | - | - | T CD8 memory 2 |
| AARS2 | S_31 | - | - | - | - | T CD8 memory 2 |
| OR7D2 | S_31 | - | - | - | - | T CD8 memory 2 |
| COX6B2 | S_31 | - | - | - | - | T CD8 memory 2 |
| ZNF665 | S_31 | - | - | - | - | T CD8 memory 2 |
| SLC12A8 | S_31 | - | - | - | Mast | T CD8 memory 2 |
| PTCHD4 | S_31 | - | - | - | - | T CD8 memory 2 |
| CMBL | S_31 | - | - | - | - | T CD8 memory 2 |
| PDP2 | S_31 | - | - | - | - | T CD8 memory 2 |
| NEK2 | S_31 | - | G2/M | - | - | T CD8 memory 2 |
| SLC14A2 | S_31 | - | - | - | - | T CD8 memory 2 |
| KCNJ5 | S_31 | - | - | - | - | T CD8 memory 2 |
| SLC24A4 | S_31 | - | - | - | - | T CD8 memory 2 |
| GUCA1B | S_31 | - | - | - | - | T CD8 memory 2 |
| PCDHB9 | S_31 | - | - | - | - | T CD8 memory 2 |
| LURAP1 | S_31 | - | - | - | - | T CD8 memory 2 |
| SEC14L4 | S_31 | - | - | - | Mast | T CD8 memory 2 |
| ZNF69 | S_31 | - | - | - | - | T CD8 memory 2 |
| CLEC4GP1 | S_31 | - | - | - | - | T CD8 memory 2 |
| CALN1 | S_31 | - | - | - | - | T CD8 memory 2 |
| MREG | S_31 | - | - | - | - | T CD8 memory 2 |
| CDKN2B-AS1 | S_31 | - | - | - | - | T CD8 memory 2 |
| CEACAM5 | S_31 | - | - | - | - | T CD8 memory 2 |
| LINC00311 | S_31 | - | - | - | Mast | T CD8 memory 2 |
| CHRNA5 | S_31 | - | - | - | - | T CD8 memory 2 |
| USP54 | S_31 | - | - | - | - | T CD8 memory 2 |
| GREB1 | S_31 | - | - | - | - | T CD8 memory 2 |
| NKPD1 | S_31 | - | - | - | - | T CD8 memory 2 |
| GRM6 | S_31 | - | - | - | - | T CD8 memory 2 |
| TM4SF1 | S_13 | Endothelial | - | - | - | CAF endothelial |
| IGFBP7 | S_13 | Endothelial ,Stroma | - | - | NK | CAF endothelial |
| SPARC | S_13 | CAF ,Stroma | - | - | - | CAF endothelial |
| CNN3 | S_13 | - | - | - | - | CAF endothelial |
| PRKCDBP | S_13 | Stroma | - | - | - | CAF endothelial |
| PRSS23 | S_13 | Stroma | - | - | NK | CAF endothelial |
| AKAP12 | S_13 | - | - | - | Mast | CAF endothelial |
| TJP1 | S_13 | - | - | - | - | CAF endothelial |
| CRIP2 | S_13 | Endothelial | - | - | - | CAF endothelial |
| MDK | S_13 | - | - | - | Plasma | CAF endothelial |
| S100A16 | S_13 | Stroma | - | - | - | CAF endothelial |
| CALD1 | S_13 | CAF | - | - | - | CAF endothelial |
| TSPAN6 | S_13 | - | - | - | - | CAF endothelial |
| SGCE | S_13 | - | - | - | - | CAF endothelial |
| FSTL1 | S_13 | CAF ,Stroma | - | - | - | CAF endothelial |
| FKBP10 | S_13 | - | - | - | - | CAF endothelial |
| PTRF | S_13 | Stroma | - | - | - | CAF endothelial |
| PLS3 | S_13 | - | - | - | - | CAF endothelial |
| WWTR1 | S_13 | - | - | - | - | CAF endothelial |
| GPRC5B | S_13 | - | - | - | - | CAF endothelial |
| SERPINH1 | S_13 | Stroma | - | - | - | CAF endothelial |
| CTTN | S_13 | - | - | - | - | CAF endothelial |
| PARVA | S_13 | - | - | - | - | CAF endothelial |
| DLC1 | S_13 | - | - | - | Mast | CAF endothelial |
| FKBP9 | S_13 | - | - | - | - | CAF endothelial |
| UBE2C | S_14 | - | G2/M ,melanoma cell cycle | Cell cycle | - | Cell cycle 1 |
| CDC20 | S_14 | - | G2/M ,melanoma cell cycle | Cell cycle | - | Cell cycle 1 |
| CCNB2 | S_14 | - | G2/M | Cell cycle | Plasma | Cell cycle 1 |
| CDKN3 | S_14 | - | melanoma cell cycle | Cell cycle | - | Cell cycle 1 |
| PLK1 | S_14 | - | - | Cell cycle | - | Cell cycle 1 |
| AURKB | S_14 | - | G2/M ,melanoma cell cycle | - | - | Cell cycle 1 |
| CCNB1 | S_14 | - | melanoma cell cycle | Cell cycle | - | Cell cycle 1 |
| DLGAP5 | S_14 | - | G2/M | Cell cycle | - | Cell cycle 1 |
| CDCA3 | S_14 | - | G2/M | - | - | Cell cycle 1 |
| NUSAP1 | S_14 | - | G2/M ,melanoma cell cycle | Cell cycle | - | Cell cycle 1 |
| BIRC5 | S_14 | - | G2/M ,melanoma cell cycle | Cell cycle | Neutrophil | Cell cycle 1 |
| TOP2A | S_14 | - | G2/M ,melanoma cell cycle | Cell cycle | - | Cell cycle 1 |
| TPX2 | S_14 | - | G2/M ,melanoma cell cycle | Cell cycle | - | Cell cycle 1 |
| RACGAP1 | S_14 | - | melanoma cell cycle | - | - | Cell cycle 1 |
| CENPF | S_14 | - | G2/M ,melanoma cell cycle | Cell cycle | - | Cell cycle 1 |
| TROAP | S_14 | - | melanoma cell cycle | - | - | Cell cycle 1 |
| CDCA8 | S_14 | - | G2/M | Cell cycle | - | Cell cycle 1 |
| PBK | S_14 | - | melanoma cell cycle | - | - | Cell cycle 1 |
| CENPE | S_14 | - | G2/M | Cell cycle | - | Cell cycle 1 |
| NUF2 | S_14 | - | G2/M ,melanoma cell cycle | - | - | Cell cycle 1 |
| BUB1B | S_14 | - | - | - | - | Cell cycle 1 |
| CDCA2 | S_14 | - | G2/M | - | - | Cell cycle 1 |
| TTK | S_14 | - | G2/M | - | - | Cell cycle 1 |
| KIF20A | S_14 | - | - | - | - | Cell cycle 1 |
| CCNA2 | S_14 | - | melanoma cell cycle | Cell cycle | - | Cell cycle 1 |
| MKI67 | S_14 | - | G2/M | Cell cycle | - | Cell cycle 1 |
| CKAP2L | S_14 | - | G2/M | Cell cycle | - | Cell cycle 1 |
| KIF15 | S_14 | - | - | Cell cycle | - | Cell cycle 1 |
| FAM64A | S_14 | - | G2/M | - | - | Cell cycle 1 |
| KIFC1 | S_14 | - | melanoma cell cycle | - | - | Cell cycle 1 |
| KIF23 | S_14 | - | G2/M | - | - | Cell cycle 1 |
| BUB1 | S_14 | - | G2/M ,melanoma cell cycle | - | - | Cell cycle 1 |
| KIF2C | S_14 | - | G2/M ,melanoma cell cycle | Cell cycle | - | Cell cycle 1 |
| CDC25C | S_14 | - | G2/M | - | - | Cell cycle 1 |
| PRC1 | S_14 | - | - | Cell cycle | - | Cell cycle 1 |
| HIST1H4F | S_14 | - | - | - | - | Cell cycle 1 |
| NCAPG | S_14 | - | - | Cell cycle | - | Cell cycle 1 |
| HMMR | S_14 | - | G2/M | Cell cycle | - | Cell cycle 1 |
| DEPDC1 | S_14 | - | - | - | - | Cell cycle 1 |
| KIF14 | S_14 | - | - | Cell cycle | - | Cell cycle 1 |
| SPAG5 | S_14 | - | - | - | - | Cell cycle 1 |
| ANLN | S_14 | - | G2/M ,melanoma cell cycle | Cell cycle | - | Cell cycle 1 |
| DEPDC1B | S_14 | - | - | - | - | Cell cycle 1 |
| HIST1H3G | S_14 | - | - | - | - | Cell cycle 1 |
| ASPM | S_14 | - | melanoma cell cycle | Cell cycle | - | Cell cycle 1 |
| HIST1H3B | S_14 | - | - | - | - | Cell cycle 1 |
| ESPL1 | S_14 | - | - | - | Mast | Cell cycle 1 |
| CCNF | S_14 | - | - | - | - | Cell cycle 1 |
| FAM72D | S_14 | - | - | - | - | Cell cycle 1 |
| KIF11 | S_14 | - | G2/M | Cell cycle | - | Cell cycle 1 |
| GAL | S_15 | - | - | - | - | Myocyte |
| KLHL41 | S_15 | - | - | - | - | Myocyte |
| BEX1 | S_15 | - | - | - | - | Myocyte |
| PPP1R27 | S_15 | - | - | - | - | Myocyte |
| NEUROD1 | S_15 | - | - | - | - | Myocyte |
| CHRNA1 | S_15 | - | - | - | - | Myocyte |
| FITM1 | S_15 | - | - | - | - | Myocyte |
| RAPSN | S_15 | - | - | - | - | Myocyte |
| HRASLS | S_15 | - | - | - | - | Myocyte |
| DES | S_15 | - | - | - | - | Myocyte |
| NTRK3 | S_15 | - | - | - | - | Myocyte |
| NEFL | S_15 | - | - | - | - | Myocyte |
| WSCD1 | S_15 | - | - | - | - | Myocyte |
| CPA1 | S_15 | - | - | - | - | Myocyte |
| MYT1L | S_15 | - | - | - | - | Myocyte |
| ATP6V1G2 | S_15 | - | - | - | - | Myocyte |
| B4GALNT1 | S_15 | - | - | - | - | Myocyte |
| ESPNL | S_15 | - | - | - | - | Myocyte |
| TMEM132E | S_15 | - | - | - | - | Myocyte |
| CKMT1A | S_15 | - | - | - | - | Myocyte |
| REEP1 | S_15 | - | - | - | - | Myocyte |
| RHOV | S_15 | - | - | - | - | Myocyte |
| CRMP1 | S_15 | - | - | - | - | Myocyte |
| PPP1R17 | S_15 | - | - | - | - | Myocyte |
| CLGN | S_15 | - | - | - | - | Myocyte |
| WFIKKN1 | S_15 | - | - | - | Plasma | Myocyte |
| POU6F2 | S_15 | - | - | - | - | Myocyte |
| KLHL25 | S_15 | - | - | - | - | Myocyte |
| FNDC5 | S_15 | - | - | - | - | Myocyte |
| IGDCC3 | S_15 | - | - | - | - | Myocyte |
| ALK | S_15 | - | - | - | - | Myocyte |
| LINC00200 | S_15 | - | - | - | - | Myocyte |
| KIF5A | S_15 | - | - | - | - | Myocyte |
| CENPV | S_15 | - | - | - | - | Myocyte |
| RNF17 | S_15 | - | - | - | - | Myocyte |
| FAM84A | S_15 | - | - | - | - | Myocyte |
| ACTN2 | S_15 | - | - | - | Mast | Myocyte |
| TLX2 | S_15 | - | - | - | - | Myocyte |
| SOX11 | S_15 | - | - | - | - | Myocyte |
| HPCA | S_15 | - | - | - | - | Myocyte |
| GABBR2 | S_15 | - | - | - | - | Myocyte |
| PYY2 | S_15 | - | - | - | - | Myocyte |
| SLC16A9 | S_15 | - | - | - | - | Myocyte |
| AGBL1 | S_15 | - | - | - | - | Myocyte |
| CLEC2L | S_15 | - | - | - | - | Myocyte |
| GKAP1 | S_15 | - | - | - | B-cell | Myocyte |
| CDH15 | S_15 | - | - | - | - | Myocyte |
| SRPK3 | S_15 | - | - | - | - | Myocyte |
| SYT7 | S_15 | - | - | - | - | Myocyte |
| CECR2 | S_15 | - | - | - | - | Myocyte |
| FN1 | S_17 | CAF | - | - | - | CAF (integrin/TGFb) |
| RCN3 | S_17 | CAF | CAF | - | - | CAF (integrin/TGFb) |
| PMP22 | S_17 | - | - | - | - | CAF (integrin/TGFb) |
| CD276 | S_17 | - | - | - | - | CAF (integrin/TGFb) |
| CYBRD1 | S_17 | CAF | CAF | - | - | CAF (integrin/TGFb) |
| VTN | S_22 | - | - | - | - | DC mature |
| ITIH1 | S_22 | - | - | - | - | DC mature |
| FBN3 | S_22 | - | - | - | - | DC mature |
| WNT5B | S_22 | - | - | - | - | DC mature |
| UPK2 | S_22 | - | - | - | - | DC mature |
| NUDT10 | S_22 | - | - | - | - | DC mature |
| ACHE | S_22 | - | - | - | - | DC mature |
| C17orf99 | S_22 | - | - | - | - | DC mature |
| GPC5 | S_22 | - | - | - | - | DC mature |
| TRIM29 | S_22 | - | - | - | - | DC mature |
| KCNH3 | S_22 | - | - | - | - | DC mature |
| GRIK4 | S_22 | - | - | - | - | DC mature |
| SLCO5A1 | S_22 | - | - | - | - | DC mature |
| MTUS2 | S_22 | - | - | - | - | DC mature |
| TMPRSS7 | S_22 | - | - | - | - | DC mature |
| VSIG8 | S_22 | - | - | - | - | DC mature |
| GPD1 | S_22 | - | - | - | - | DC mature |
| ANXA10 | S_22 | - | - | - | - | DC mature |
| CD40LG | S_19 | T CD4 | - | - | T cell | T CD4 memory/naive |
| TNFSF8 | S_19 | T CD4 | - | - | T cell | T CD4 memory/naive |
| IL7R | S_19 | T CD4 ,T CD4 naive ,T CD8 naive | - | - | T cell | T CD4 memory/naive |
| CAMK4 | S_19 | T CD4 ,T CD4 naive ,T CD8 naive | - | - | T cell | T CD4 memory/naive |
| PRKCQ-AS1 | S_19 | T CD8 naive | - | - | NK | T CD4 memory/naive |
| FLT3LG | S_19 | T CD4 | - | - | - | T CD4 memory/naive |
| TCF7 | S_19 | T CD4 ,T CD4 naive ,T CD8 naive | T- cell | - | T cell | T CD4 memory/naive |
| GINS2 | S_28 | - | G1/S ,melanoma cell cycle | - | - | Cell cycle 2 |
| CDC45 | S_28 | - | G1/S ,melanoma cell cycle | - | - | Cell cycle 2 |
| RFC5 | S_28 | - | melanoma cell cycle | - | - | Cell cycle 2 |
| WDHD1 | S_28 | - | - | - | - | Cell cycle 2 |
| MCM2 | S_28 | - | G1/S ,melanoma cell cycle | - | - | Cell cycle 2 |
| DTL | S_28 | - | G1/S ,melanoma cell cycle | - | - | Cell cycle 2 |
| E2F1 | S_28 | - | - | - | - | Cell cycle 2 |
| POLE2 | S_28 | - | - | - | - | Cell cycle 2 |
| RMI2 | S_28 | - | - | - | pDC | Cell cycle 2 |
| POLA1 | S_28 | - | G1/S | - | - | Cell cycle 2 |
| MCM10 | S_28 | - | - | - | pDC | Cell cycle 2 |
| HELLS | S_28 | - | G1/S | - | - | Cell cycle 2 |
| CDC6 | S_28 | - | G1/S ,melanoma cell cycle | - | - | Cell cycle 2 |
| WDR76 | S_28 | - | G1/S | - | - | Cell cycle 2 |
| MYH1 | S_23 | - | - | - | - | T CD8 memory 1 |
| MYLK2 | S_23 | - | - | - | - | T CD8 memory 1 |
| ASB5 | S_23 | - | - | - | - | T CD8 memory 1 |
| MYBPH | S_23 | - | - | - | - | T CD8 memory 1 |
| DDN | S_23 | - | - | - | - | T CD8 memory 1 |
| DUSP27 | S_23 | - | - | - | - | T CD8 memory 1 |
| BEST3 | S_23 | - | - | - | Plasma | T CD8 memory 1 |
| TNFRSF25 | S_26 | - | - | - | T cell | T CD4 reg |
| FAAH2 | S_26 | T CD4 | - | - | - | T CD4 reg |
| CCR4 | S_26 | T CD4 ,T CD4 Treg | - | - | - | T CD4 reg |
| FBLN7 | S_26 | T CD4 | - | - | T cell | T CD4 reg |
| PBX4 | S_26 | T CD4 | - | - | T cell | T CD4 reg |
| RORA | S_26 | T CD4 Treg | - | - | - | T CD4 reg |
| PBXIP1 | S_26 | T CD4 | - | - | T cell | T CD4 reg |
| GIMAP5 | S_29 | T CD4 naive | - | - | T cell | Lymphoid cells |
| GIMAP7 | S_29 | - | - | - | T cell | Lymphoid cells |
| GIMAP4 | S_29 | - | - | - | T cell | Lymphoid cells |
| GIMAP6 | S_29 | - | - | - | NK | Lymphoid cells |
| FYB | S_29 | T CD4 ,T-cell | - | - | - | Lymphoid cells |
| ACOT7 | S_30 | Malignant | - | - | - | Cell cycle/dna repair |
| RAD51C | S_30 | Malignant | - | - | - | Cell cycle/dna repair |
| TMEM106C | S_30 | - | - | - | - | Cell cycle/dna repair |
| C19orf48 | S_30 | - | - | - | - | Cell cycle/dna repair |
| STMN1 | S_30 | Malignant | - | Cell cycle | pDC | Cell cycle/dna repair |
| DHCR24 | S_30 | - | - | - | - | Cell cycle/dna repair |
| DTYMK | S_30 | - | melanoma cell cycle | - | - | Cell cycle/dna repair |
| WDR34 | S_30 | - | melanoma cell cycle | - | - | Cell cycle/dna repair |
| TMEM14A | S_30 | - | - | - | - | Cell cycle/dna repair |
| CTNNAL1 | S_30 | - | - | - | - | Cell cycle/dna repair |
| CDK2 | S_30 | Malignant | melanoma | - | - | Cell cycle/dna repair |
| ALG14 | S_33 | - | - | - | - | Plasma (identical protein binding) |
| PYCR1 | S_33 | - | - | - | Plasma | Plasma (identical protein binding) |
| CAV1 | S_33 | - | - | - | Plasma | Plasma (identical protein binding) |
| MAGED1 | S_33 | - | - | - | pDC | Plasma (identical protein binding) |
| CHPF | S_33 | - | - | - | Plasma | Plasma (identical protein binding) |
