## Supplementary figures and images for "Immune cell type signature discovery and random forest classification for analysis of single cell gene expression datasets"

### SuppFig1.pdf

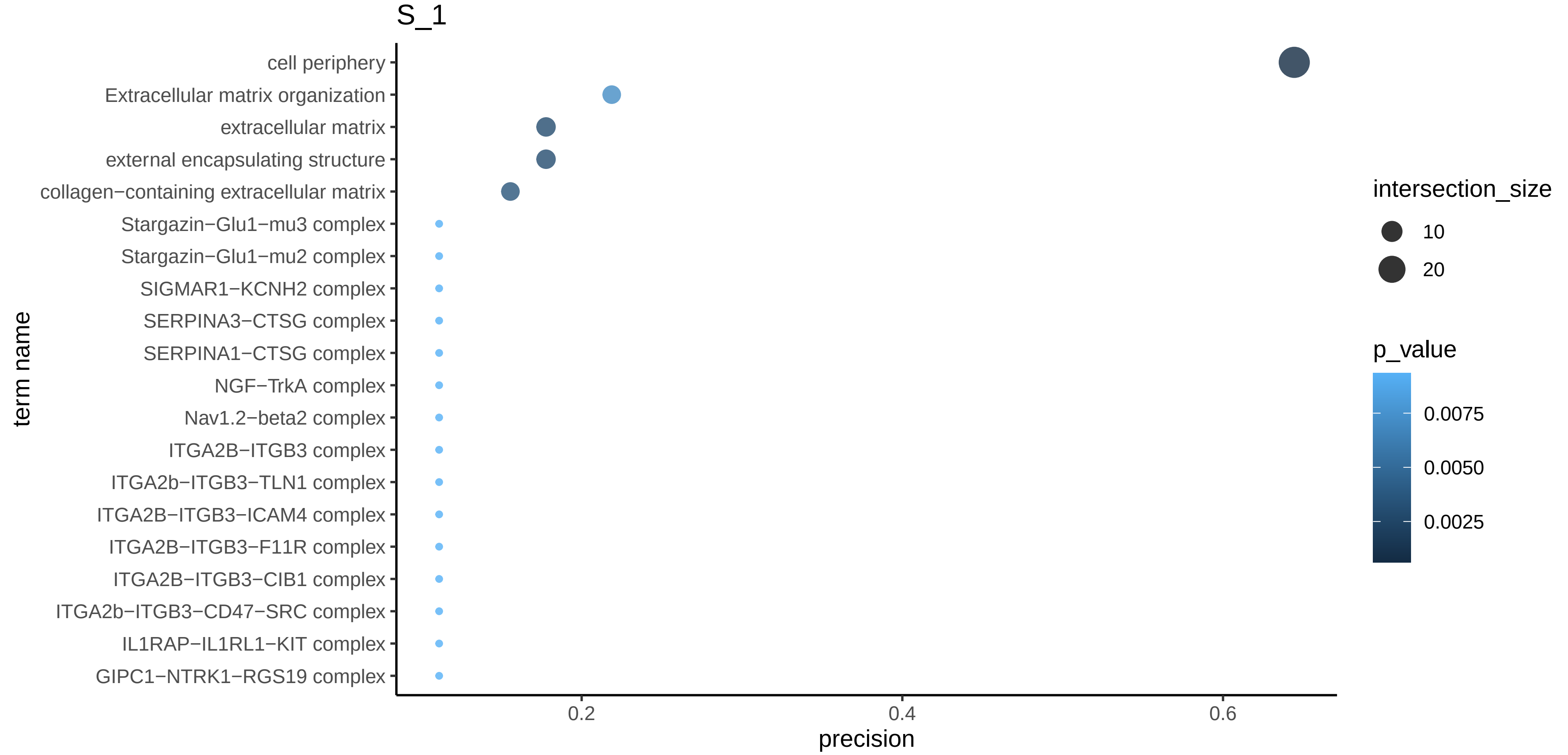

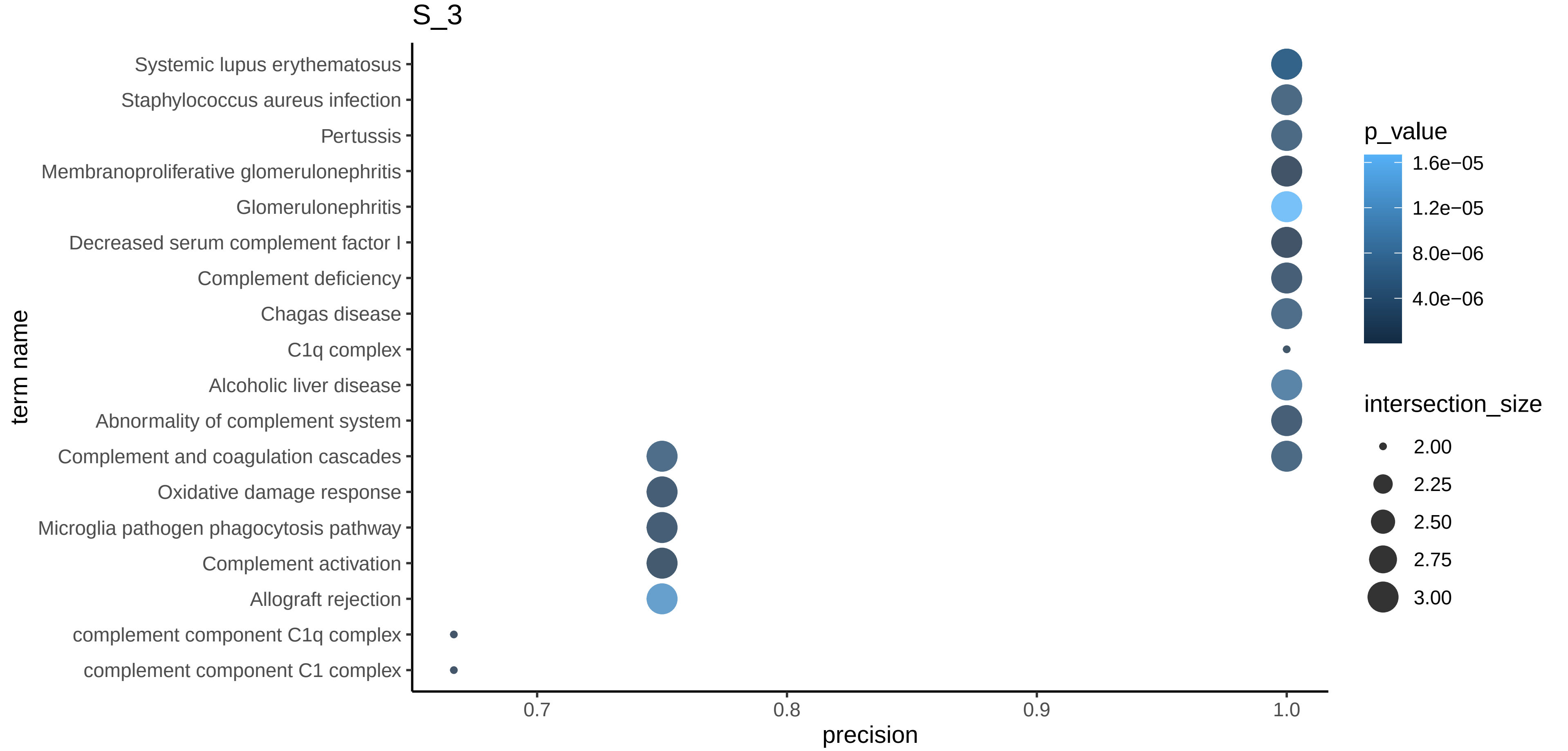

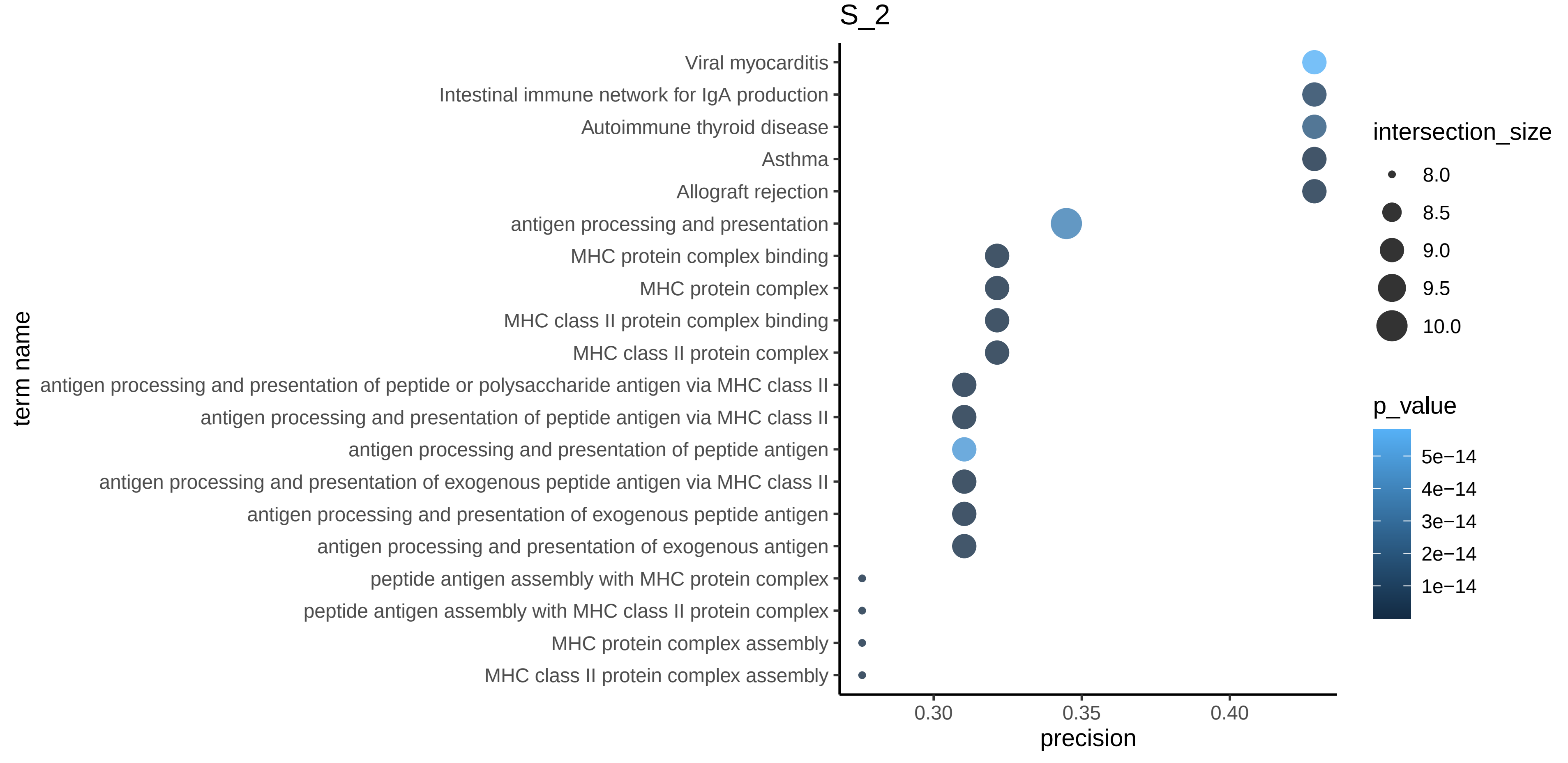

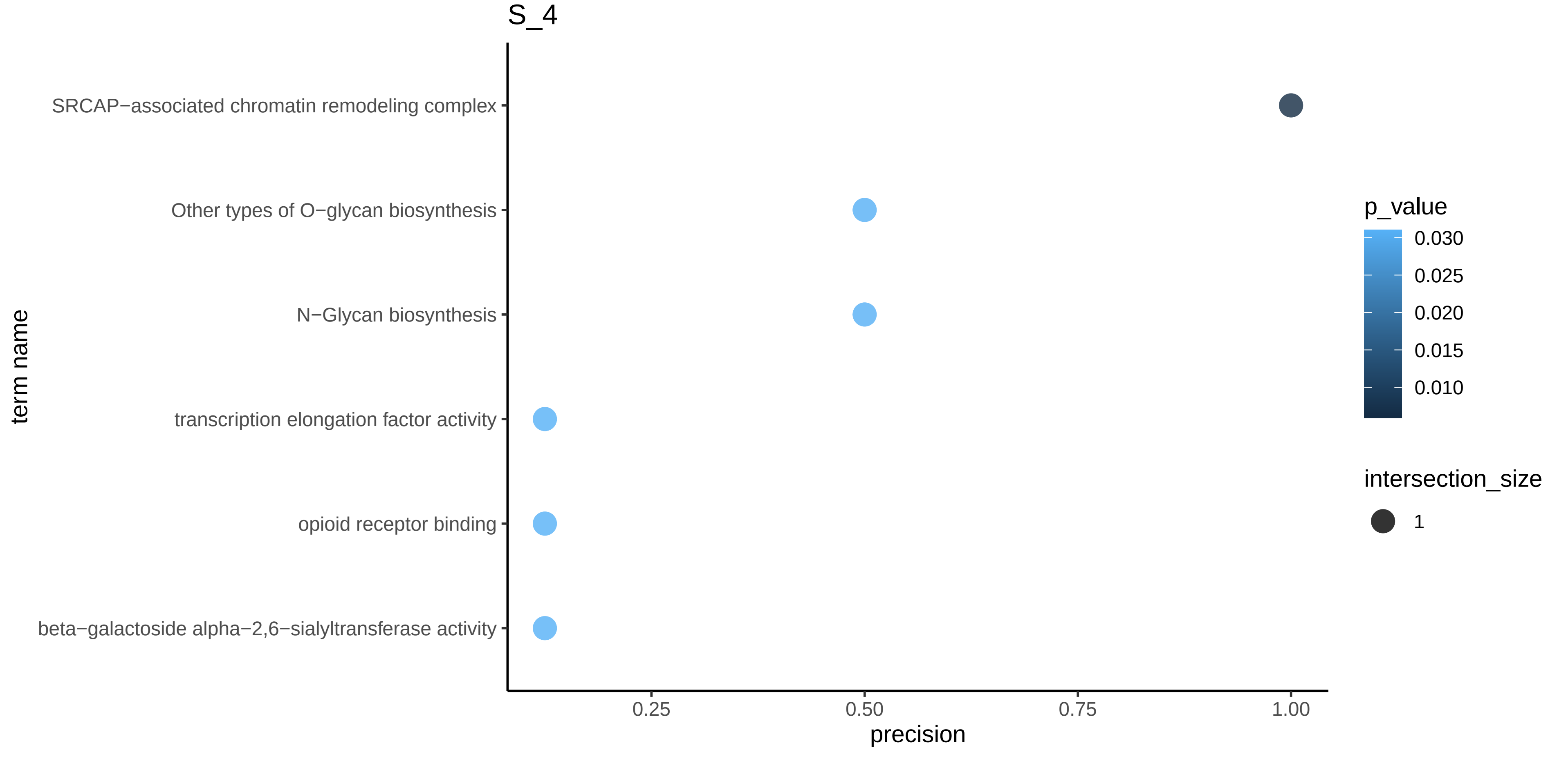

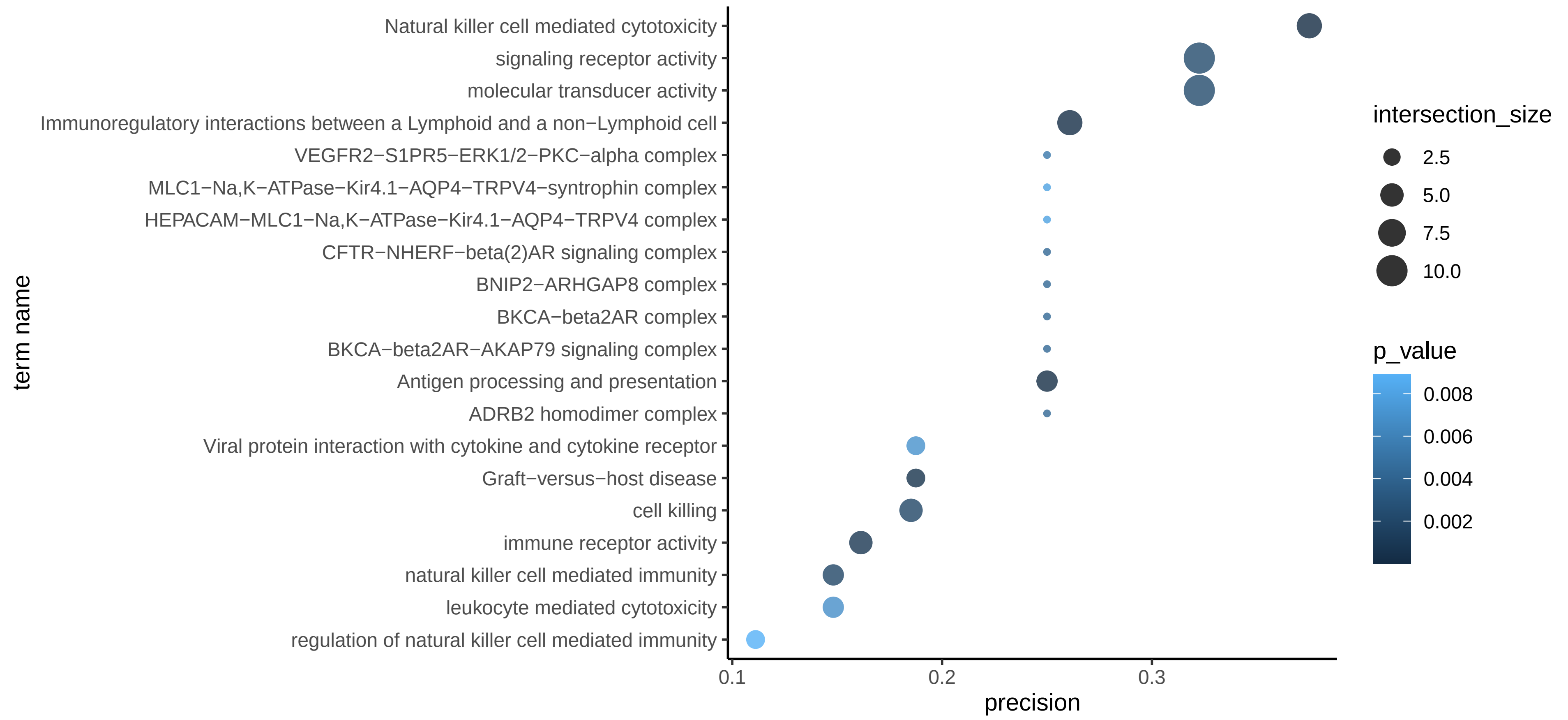

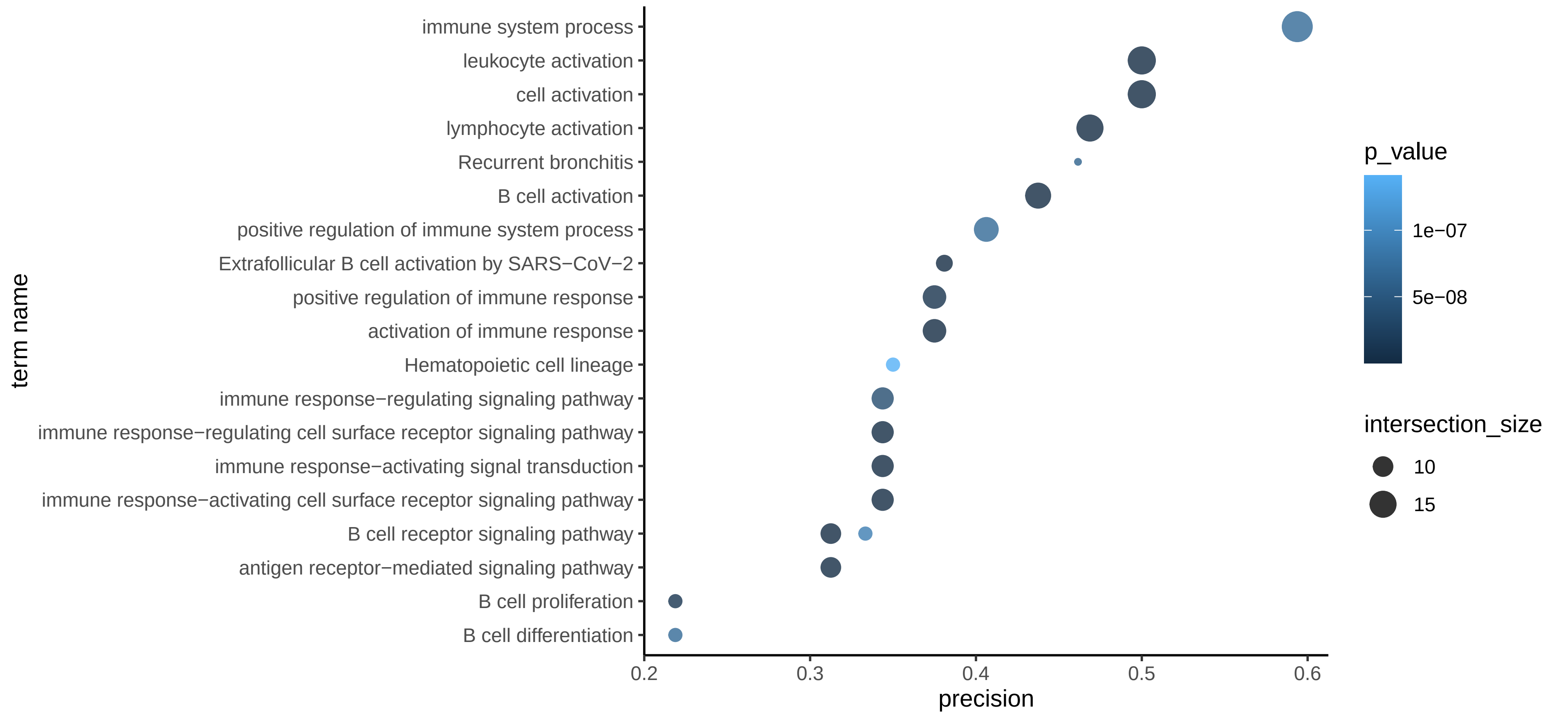

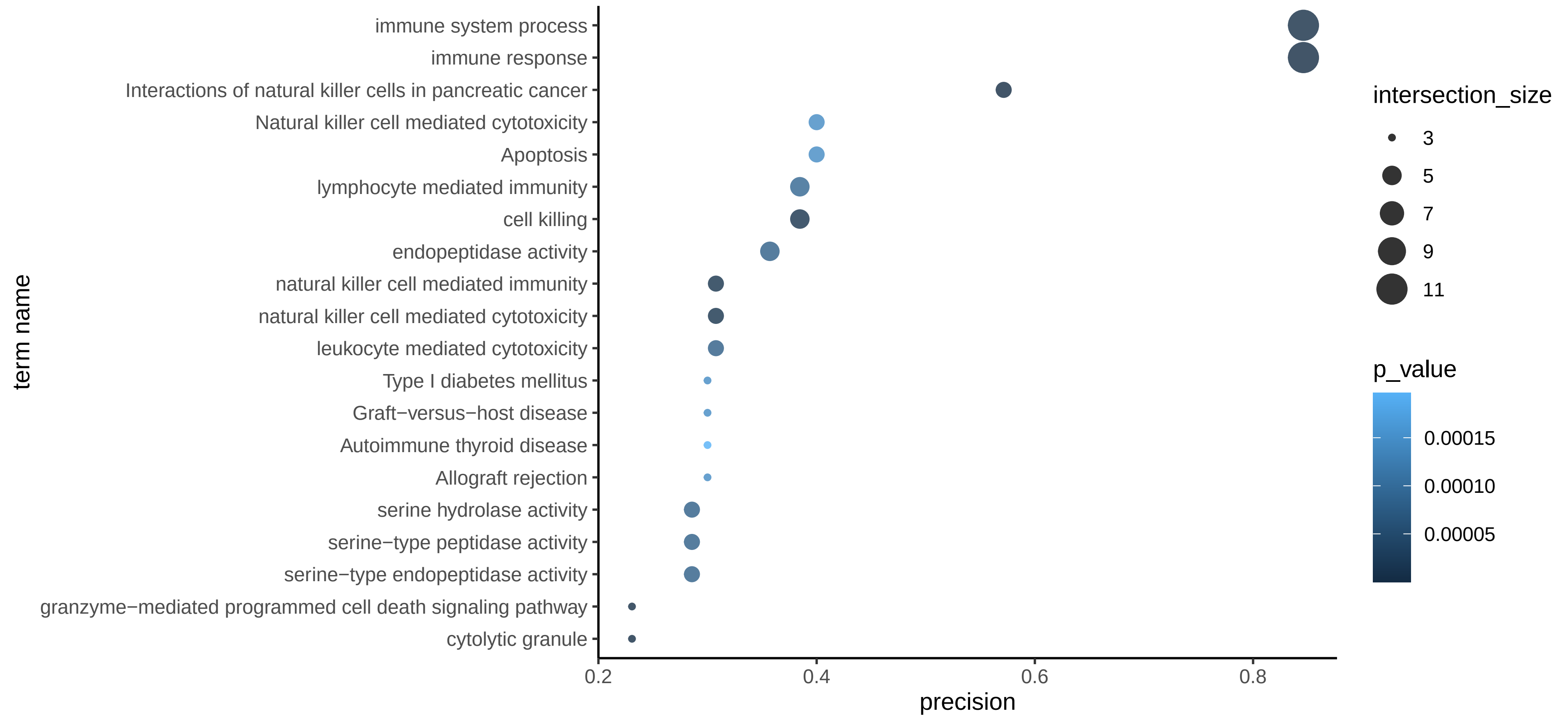

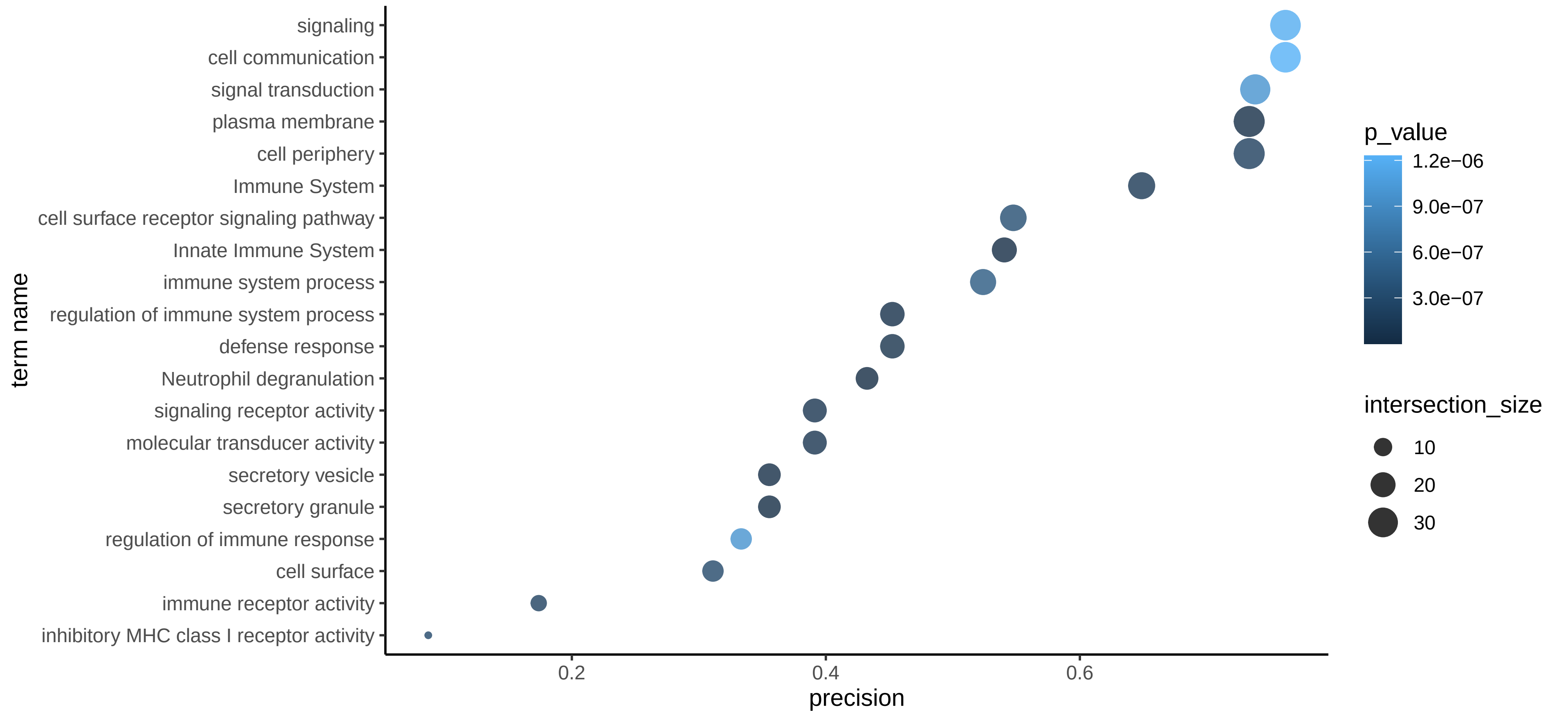

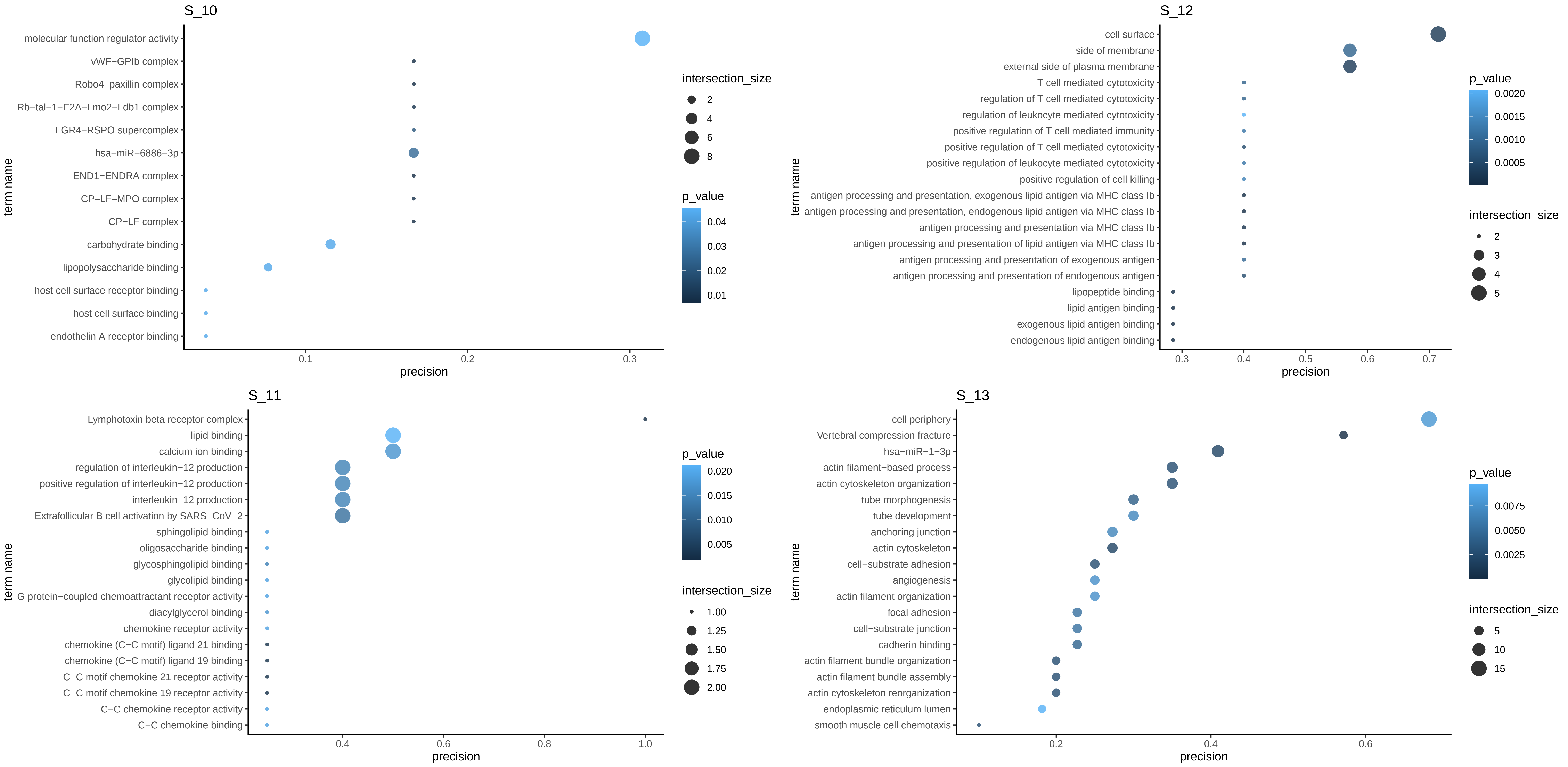

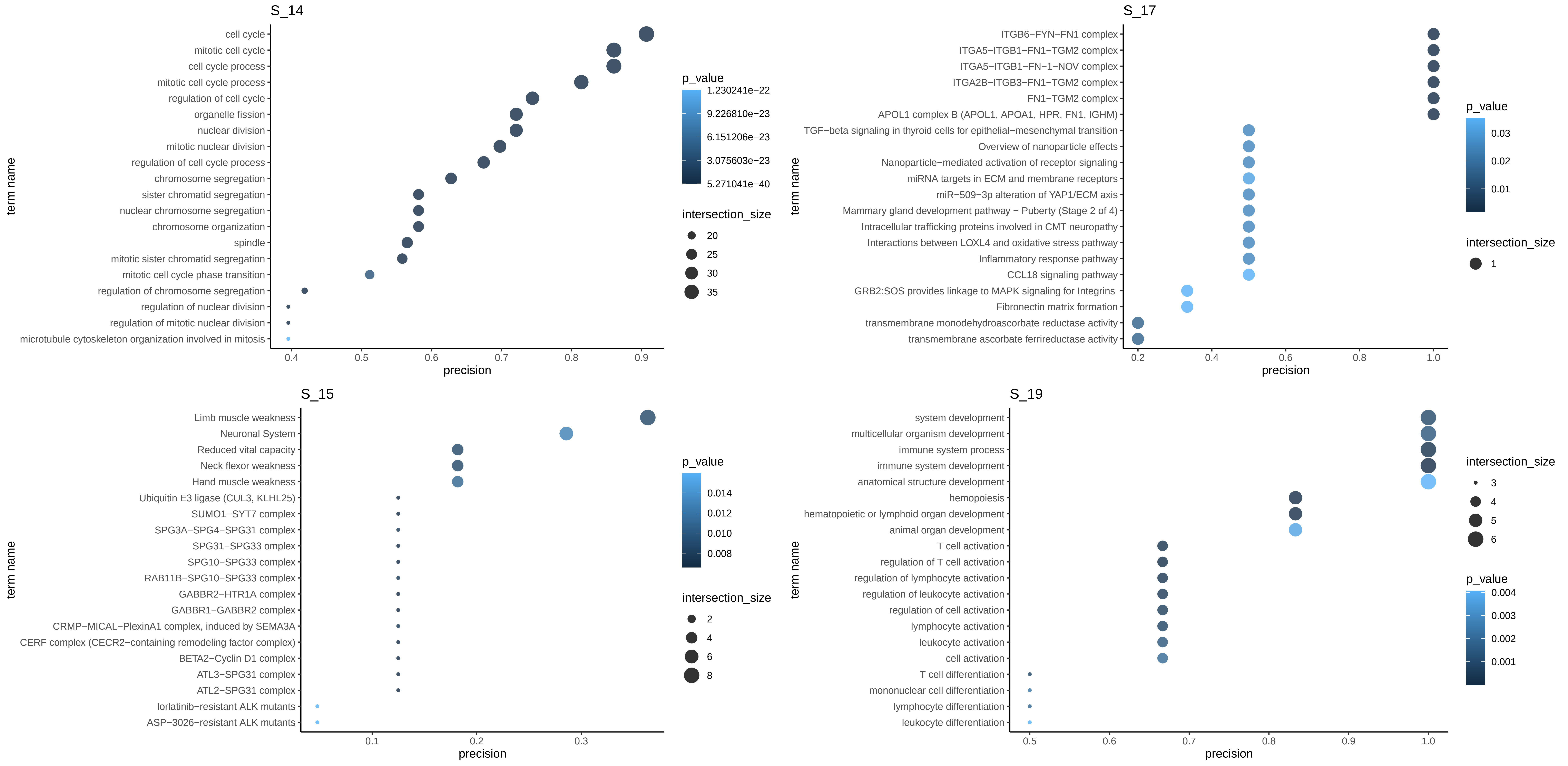

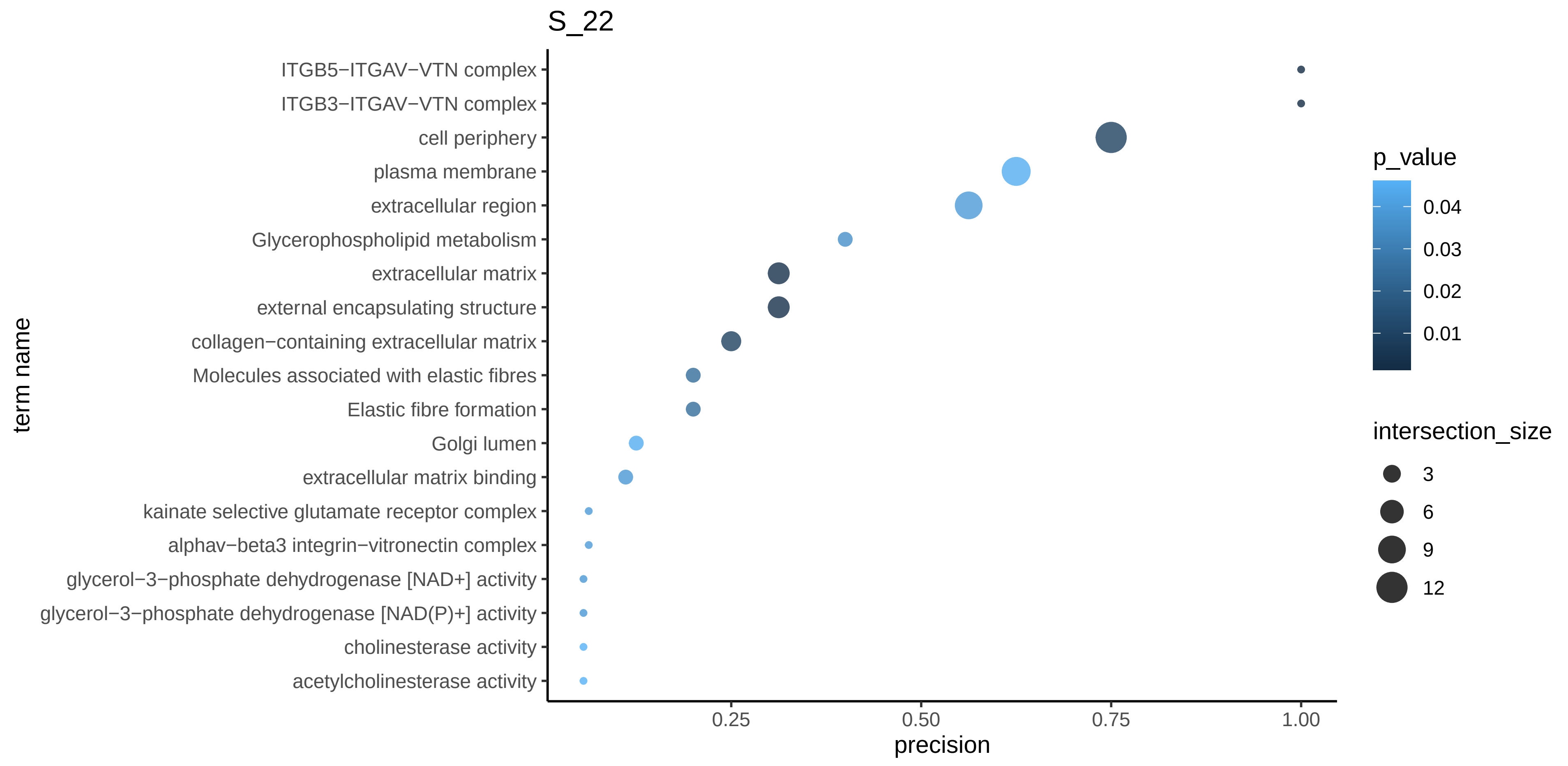

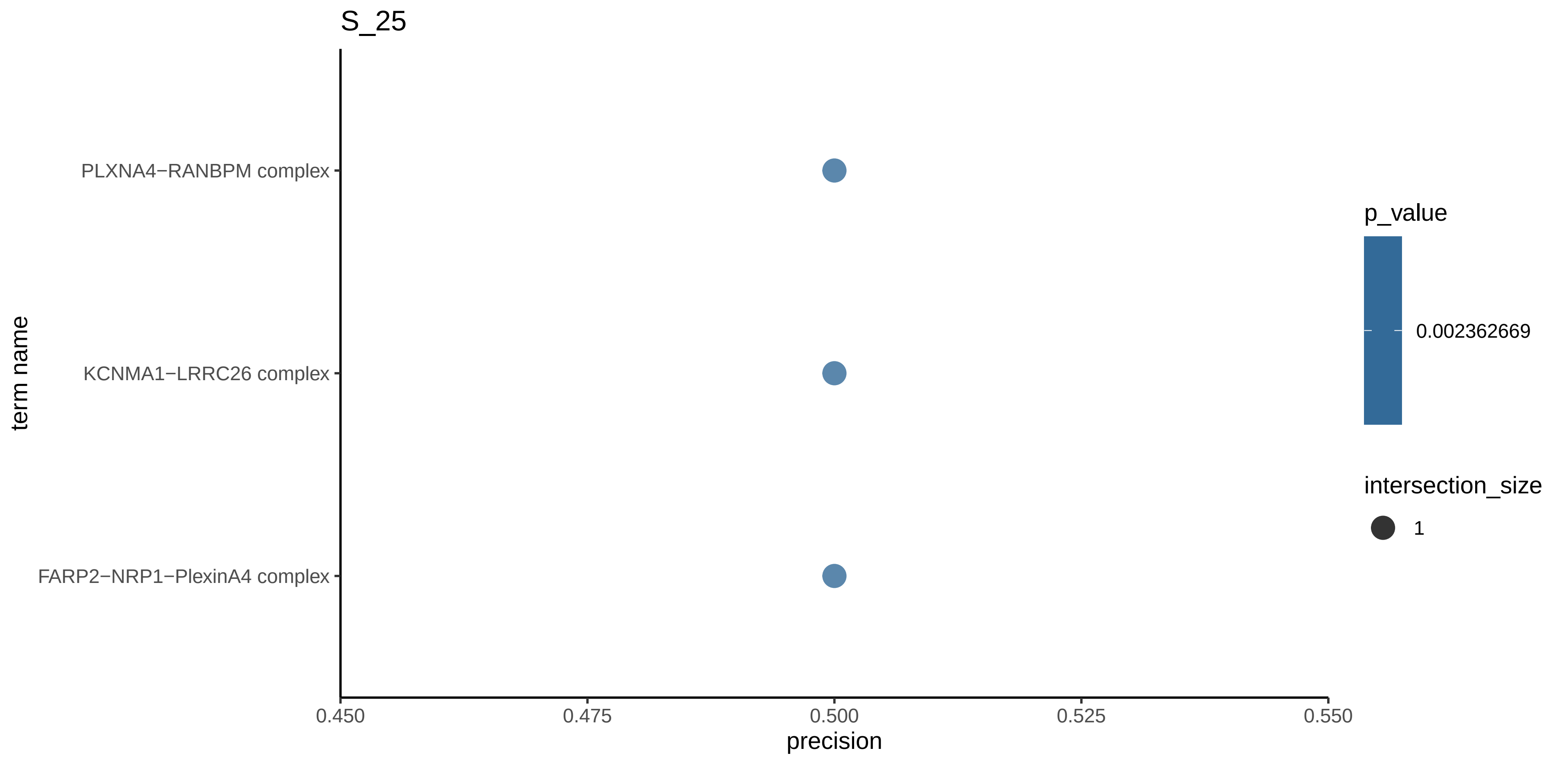

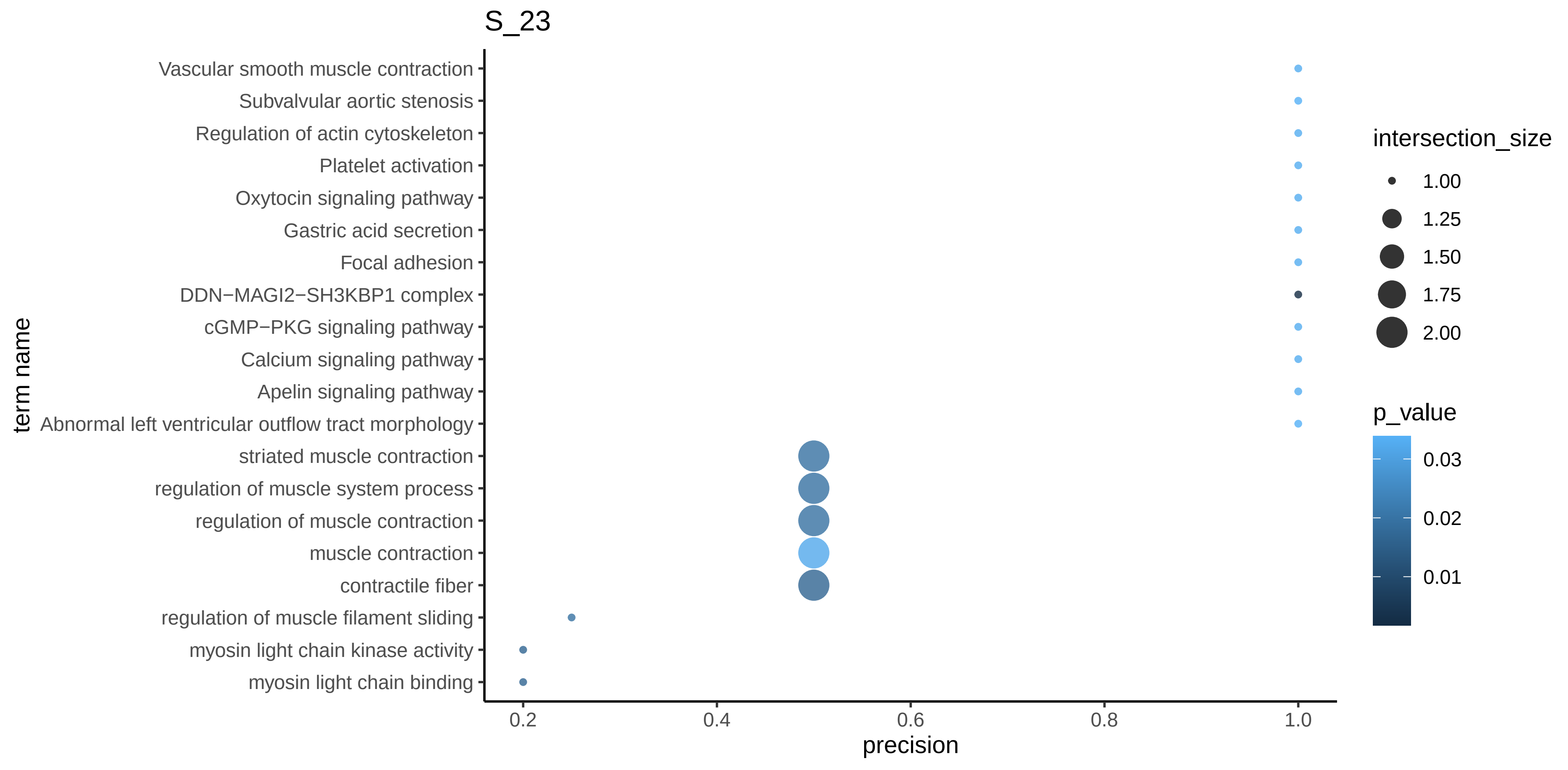

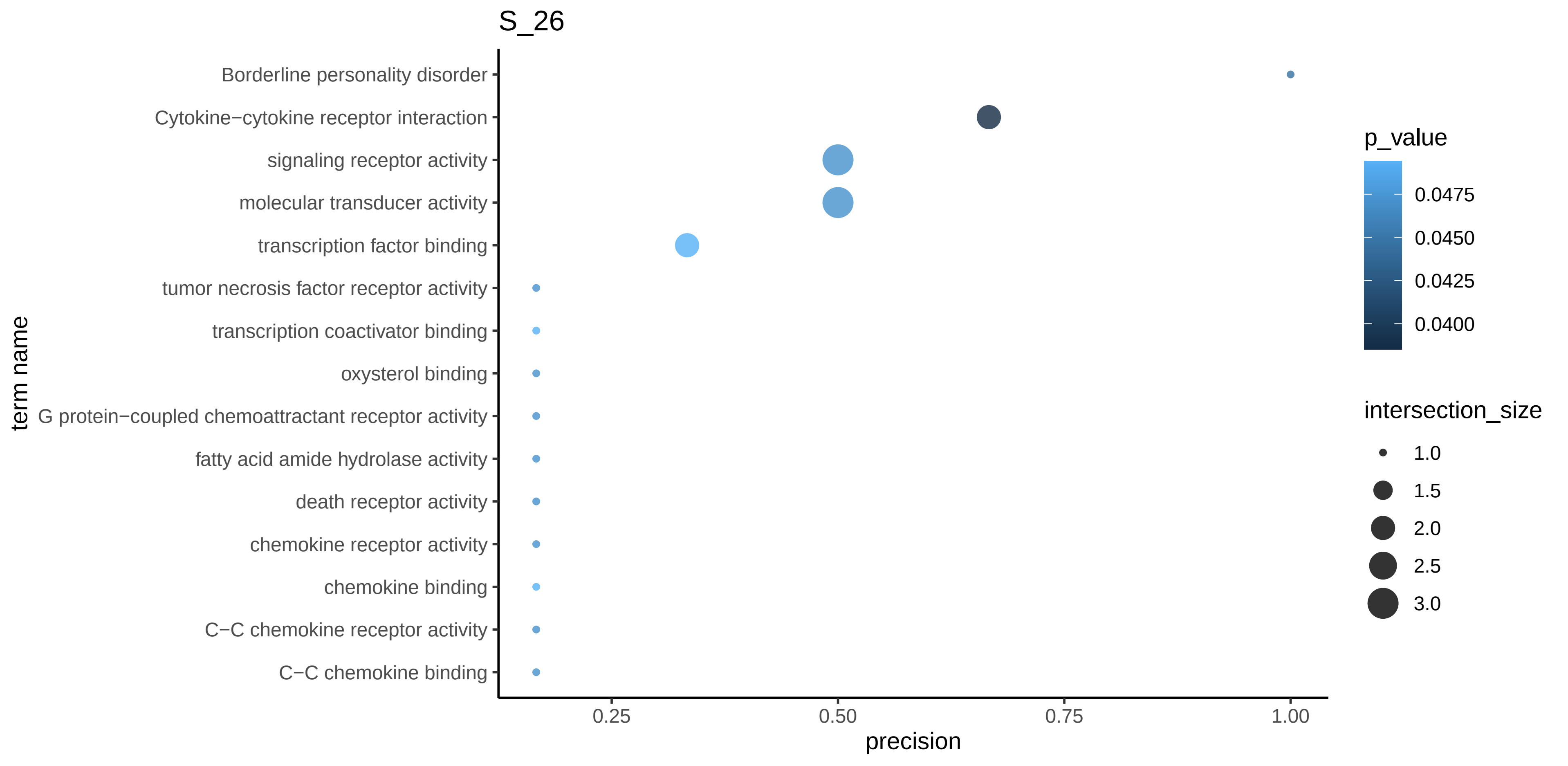

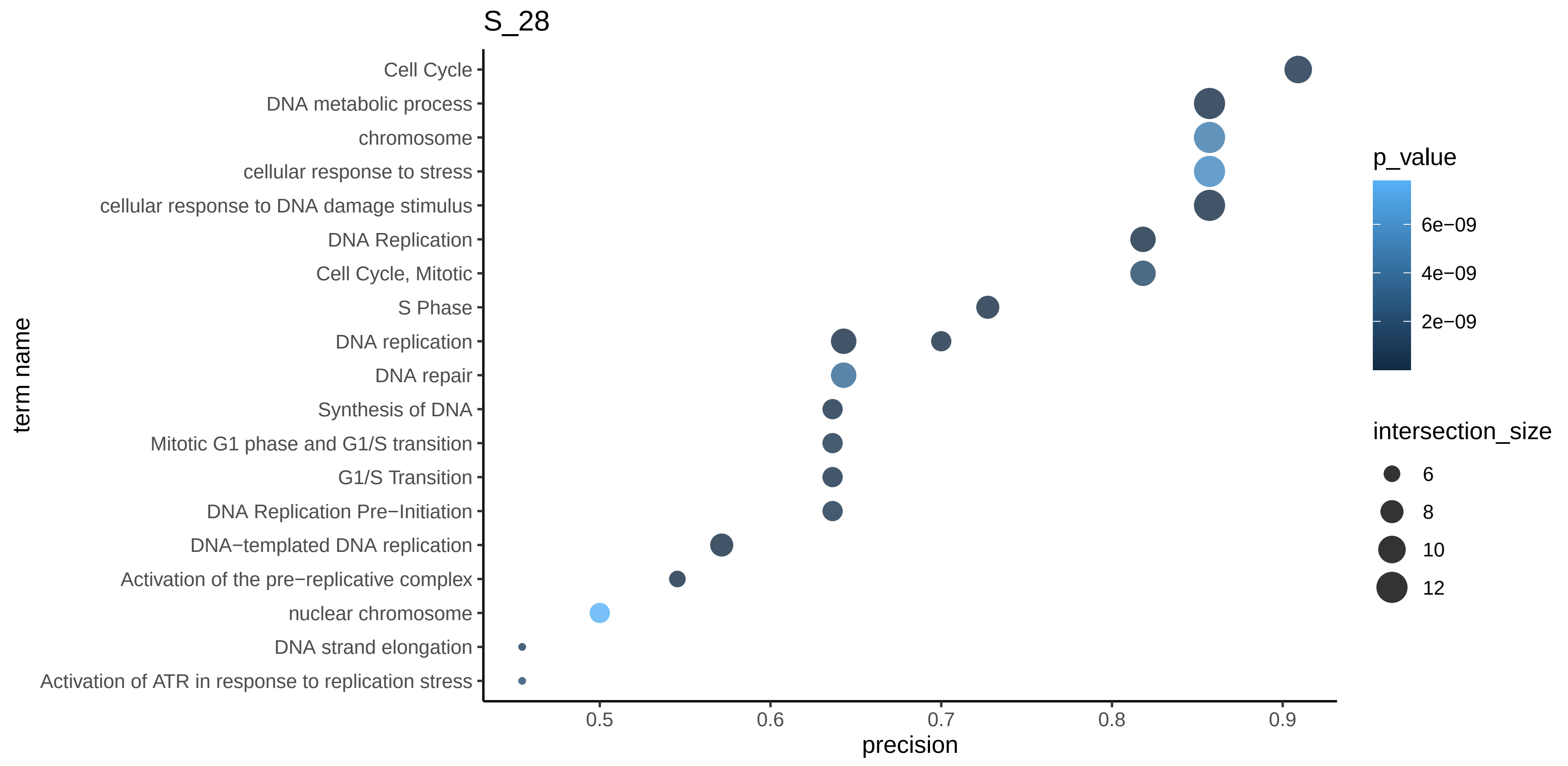

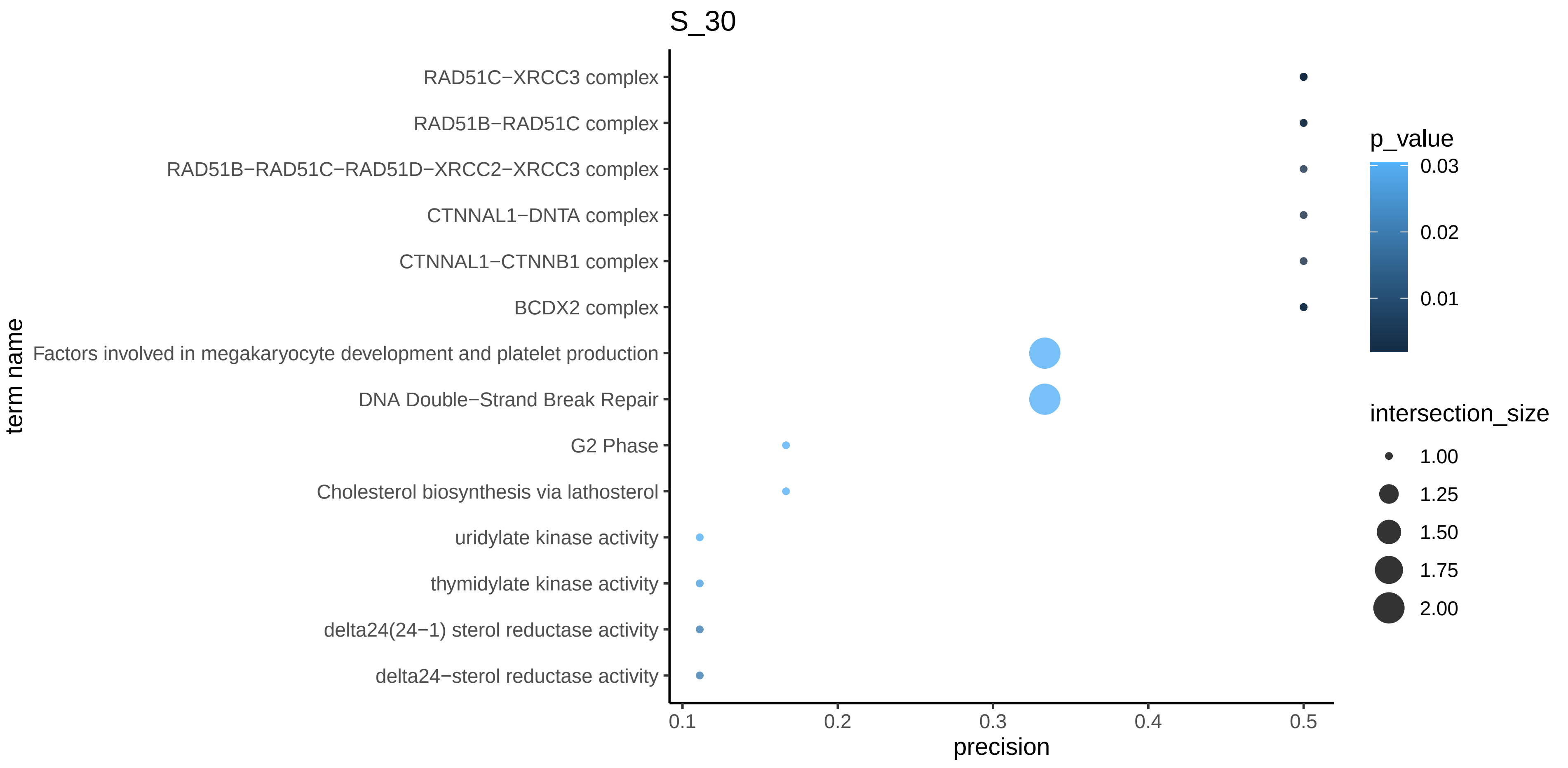

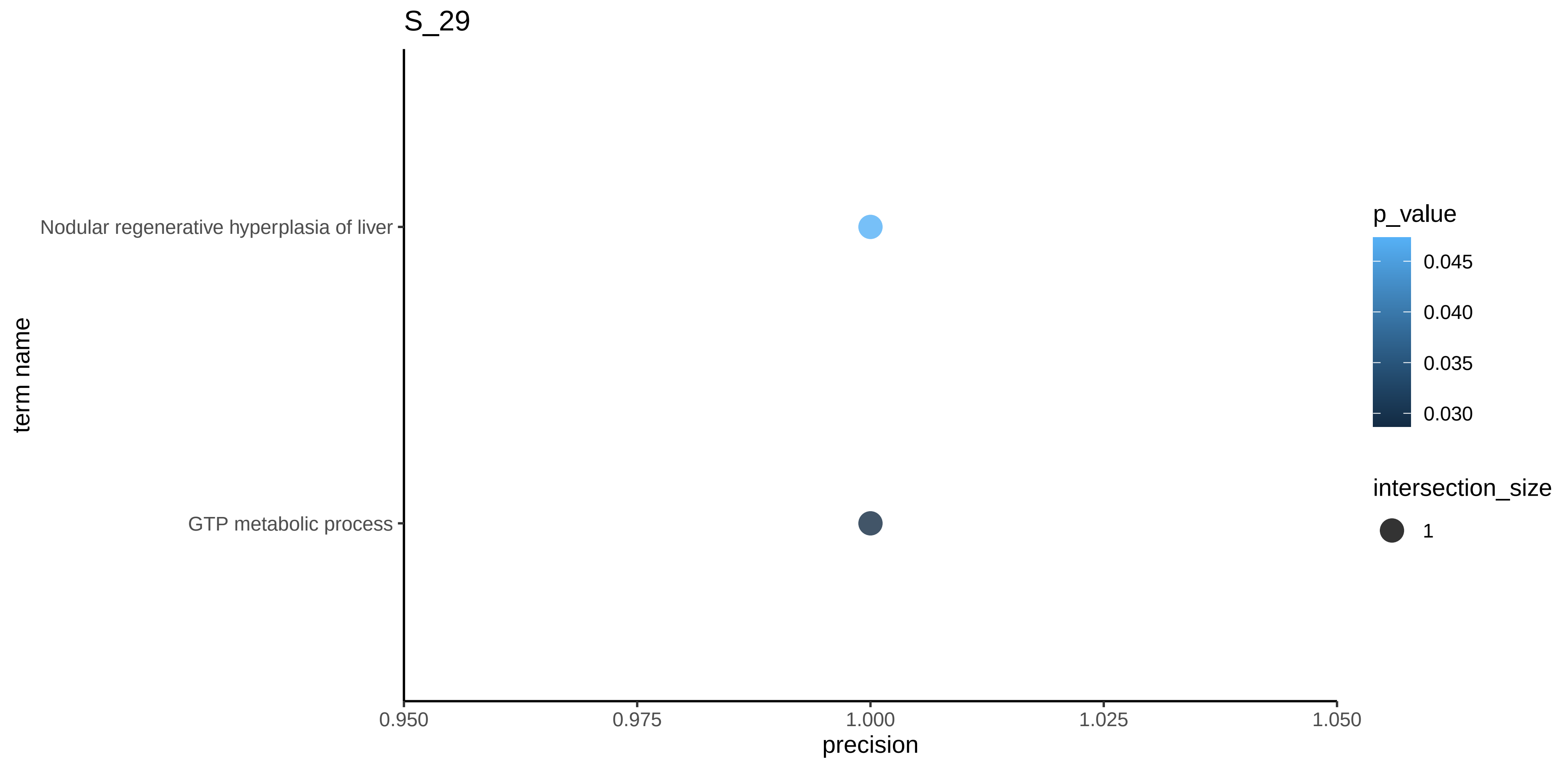

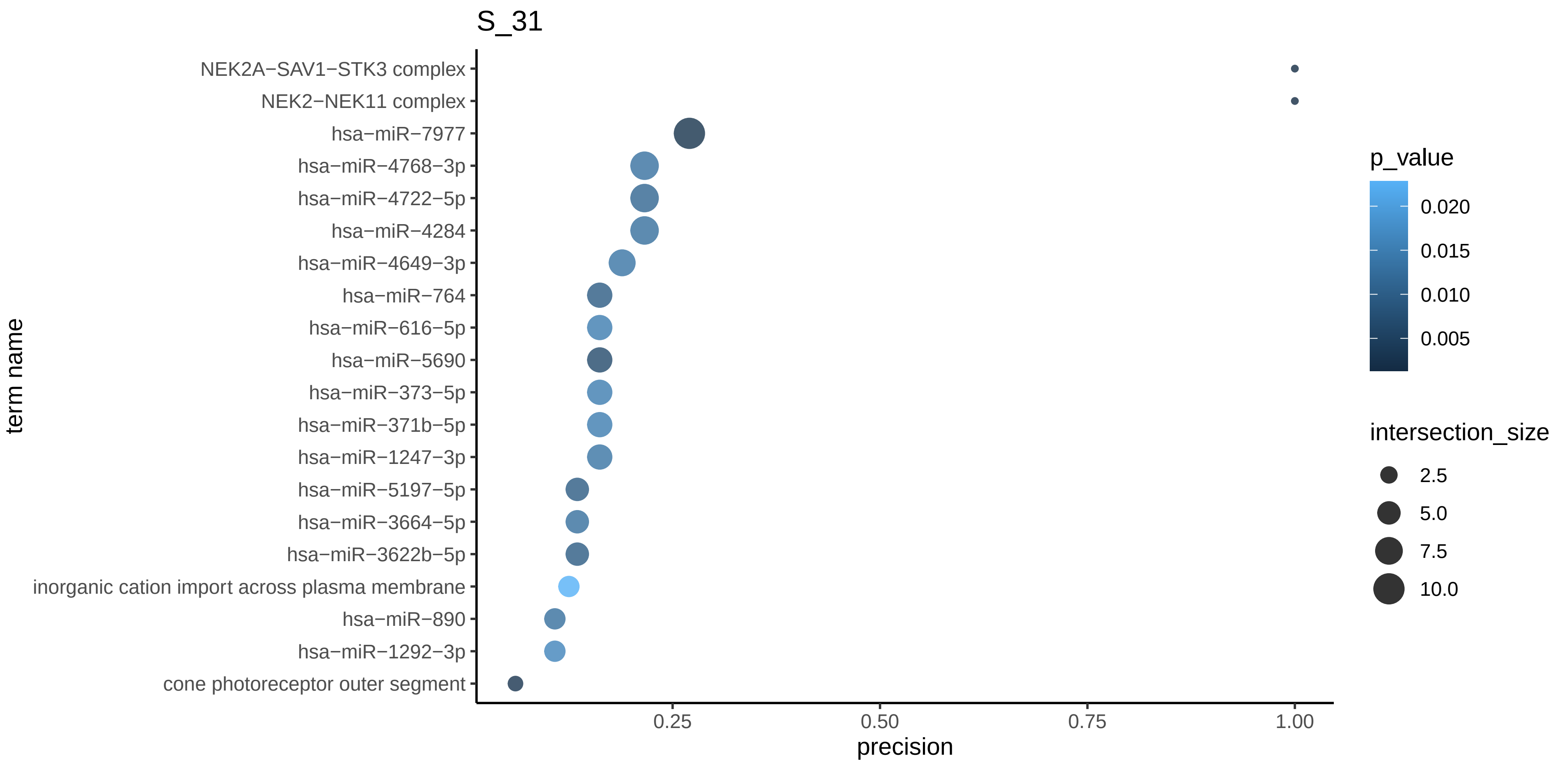

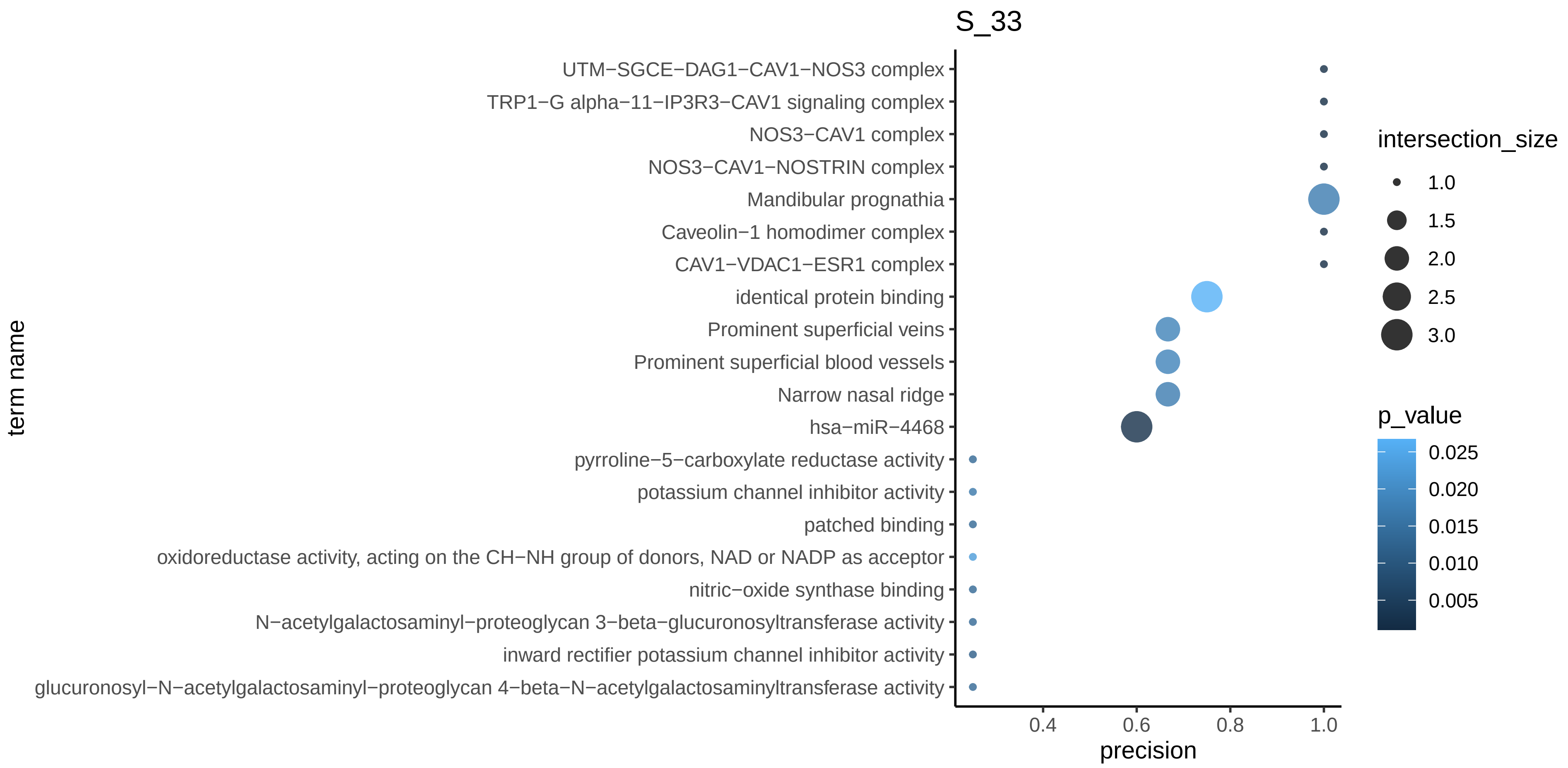
